# Bioinformatic screening of genes present only in well regenerating vertebrates reveals novel FGF and purinergic signaling modulator - *c-Answer*

**DOI:** 10.1101/494609

**Authors:** Daria D. Korotkova, Vassily A. Lyubetsky, Anastasia S. Ivanova, Lev I. Rubanov, Alexander V. Seliverstov, Oleg A. Zverkov, Natalia Yu. Martynova, Alexey M. Nesterenko, Maria B. Tereshina, Leonid Peshkin, Andrey G. Zaraisky

## Abstract

The genetic basis of higher regenerative capacity of fishes, amphibians and reptiles compared to birds and mammals is still poorly understood. Though it is thought to be a result of restructuring in the regulatory network of a static set of genes, we argued that it could be due to the loss of genes essential for regeneration. In the present work, we formulate a bioinformatic approach to systematic search for the such genes. Having identified them, we further investigated one we dubbed “*c-Answer*”, which encodes a membrane protein, regulating the regeneration of body appendages and the telencephalic development through binding to FGFR and P2Y1 receptors and promoting MAPK/ERK and purinergic signaling. The obtained data suggest that elimination of *c-Answer* in the ancestors of warm-blooded animals conditioned the decreased activity of at least two signaling pathways, which in turn could contribute to changes in mechanisms that regulate the forebrain development and regeneration.

## INTRODUCTION

Restructuring of the cis-regulatory elements of the gene network, which consist of approximately the same set of genes, is thought to underlay most of the evolutionary events, in particular, the reduction of the appendage regenerative capacity in birds and mammals (warm-blooded animals) comparing to well regenerating fishes, amphibians and reptiles (cold-blooded animals) (Rodríguez-Trelles et al., 2003; Wray, 2007). However, we have recently demonstrated that genes encoding thioredoxin domain-containing secreted protein Ag1 and small GTPases Ras-dva1 and Ras-dva2, which are essential for the regeneration of body appendages in fishes and frogs, were eliminated in evolution of birds and mammals (Ivanova et al., 2013a, 2015; Tereshina et al., 2014). Accordingly, this allowed us to hypothesize that, at least in part, low regenerative capacity of warm-blooded animals could be explained by extinction of *Ag1* and *Ras-dva1/2* in evolution. In turn, assuming these data, one may suppose that there could also be other genes that might be lost during evolution of birds and mammals, however still be involved in regulation of regeneration in well regenerating cold-blooded vertebrates. In the present work, we proposed a bioinformatic method (algorithm and computer program) for systematic search for such genes combined with the experimental testing of the predicted genes involvement in the *Xenopus laevis* tadpoles tail and hindlimb bud regeneration.

By using the developed bioinformatic approach, we identified several genes missing in warm-blooded animals and selected ones demonstrating an increased expression during regeneration of the amputated tadpole tail and hindlimb bud. Basing on the protein sequence analysis, we selected one gene that encodes previously unknown putative membrane protein and studied its functions in depth. As we have demonstrated, this gene is expressed predominantly in the presumptive neural plate beginning from the late gastrula stage and it is sharply activated in cells of the wound epithelium at the 1^st^ day after amputation of the tadpole tail and hindlimb bud. Down-regulation of this gene by anti-sense morpholino and CRISPR/Cas9 resulted in diminishing of the overall tadpole size, specifically eye size, and a retardation of tadpole tail regeneration. On the other hand, overexpression of the identified gene elicited reverse effects, i.e. an increase of the telencephalon, eyes, including ectopic eye differentiation, and restoration of tail regeneration in the ‘refractory’ period when the tail normally cannot regenerate. We have also shown that the membrane protein encoded by this gene can bind two types of receptors involved in signaling that regulate the development of the telencephalon, eyes and regeneration: the Fgf receptors, FGFR1-4, and the receptor of extracellular ADP, P2Y1. Accordingly, we named this protein c-Answer after <u>c</u>old-blooded <u>An</u>imal <u>s</u>pecific <u>w</u>ound <u>e</u>pithelium receptor-binding protein. Together with our previous data on *Ag1* and *Ras-dva* genes, the obtained results confirm that such significant evolutionary changes as the loss of ability to regenerate major body appendages along with the progressive evolution of the forebrain, which are characteristic to warm-blooded animals, could be caused by the loss in their ancestors of a gene set that regulates regeneration and brain development in cold-blooded species.

## RESULTS

### Genes missing in warm-blooded vertebrates identified by our bioinformatic approach

We developed an approach aimed at identifying genes present in cold-blooded animals that have no orthologs, i.e. direct homologs, in warm-blooded species (Lyubetsky et al., 2017; Zverkov et al., 2015). For the purposes of our study, we regarded orthologs as a pair of homologs in distinct species which keep local genomic synteny, i.e. have at least one pair of independent homologous genes in their vicinity. The assumption being that if a given cold-blooded animal gene was lost during evolution in warm-blooded animals, its intra-chromosome neighbors in cold-blooded animals should not have homologs in warm-blooded animals in the vicinity of any homolog of the lost gene. Homology was established using protein sequences.

To establish orthology, we used the two-step strategy. The genome of *Xenopus tropicalis*, which presumably contains a full collection of genes present in cold-blooded animals and is well-sequenced, was used as a reference, or basic species. At the first step, by using previously developed algorithm (called ClusterZSL) (Lyubetsky et al., 2013; Rubanov et al., 2016; Zverkov et al., 2012, 2015) for each gene of *X. tropicalis* we formed a cluster of the most homologous genes from the selected representatives of cold- and warm-blooded species with well sequenced and annotated genomes. Several orthology inference methods including the local synteny consideration were compared previously using five mammalian genomes (Jun et al., 2009a).

At the second step, within each cluster we chose homologs, which would keep local genomic synteny in cold-blooded species, but would not in warm-blooded ones. These homologs were considered as genes lost in evolution by warm-blooded animals.

The developed algorithm can operate on any two juxtaposed groups of animals, e.g. anamniotes vs. amniotes, apodes vs. tetrapodes, short vs. long lifespan etc., that henceforth are referred to as “lower” and “upper” sets. These groups can be generated basing on any trait that is present in lower species but is not in upper ones. In our study “lower” set are well regenerating cold-blooded animals (fish, amphibians, reptiles) while “upper” are poorly regenerating warm-blooded ones (birds, mammals). In turn, sets are subdivided into parts. By this term we designated a group of selected species belonging to some biological class of vertebrates.

Numeric parameters *p*_i_ and *q_j_* were assigned to each upper and lower part, respectively. Then, in the basic species (*X. tropicalis*) we tried to identify the genes with disrupted synteny in at least *p_i_* upper species of the *i*-th part and undisrupted in at least *q_j_* lower species of the *j*-th part; given this, some upper and lower species would lack or have the gene, respectively. In an extreme case, *p_i_* equals the number of upper species and *q_j_* = 1 for all *i* and *j*. A decreasing number of gene paralogs was used as an additional condition with a numeric parameter *r_j_*. Specifically, if the number of paralogs of a gene in the basic species exceeds the number of its paralogs in an upper species in accordance with the specified parameter *r_j_*≥1, the gene is considered lost.

All protein-coding genes from the basic species were tested using the 2- and 3-species modes. In the 2-species mode, homologs *X\** of gene *X* in each lower species were checked for synteny. In this mode, the following two conditions were verified for to establish that a given gene has no orthologs in the upper species. (1) A pair of genes *Y* and *Z* are defined in the basic species as different from *X* and each other and co-localized within a window of the size 2*l*; their homologs *X*\*, *Y*\*, *Z*\* must co-localize within windows of the size 2*l*_1_ in lower species as shown by bold arrows (Figure 1A). (2) There are no homologs *X*\*\* in upper species or their synteny is disrupted. The latter means that no homolog of any gene *S* within the window in the basic species exists within windows of the size 2*l*_2_ among upper species (Figure 1A). The genes *Y, Z, S* will be referred to as witnesses. The algorithm parameters specify the desired number of witnesses. 2Mbp was chosen as the value of *1=1_i_=1_2_*, basing on our empirical observation that at least one witness could be always found at the distance of 2Mbp or less for all well-established orthologs.

**Figure 1.**
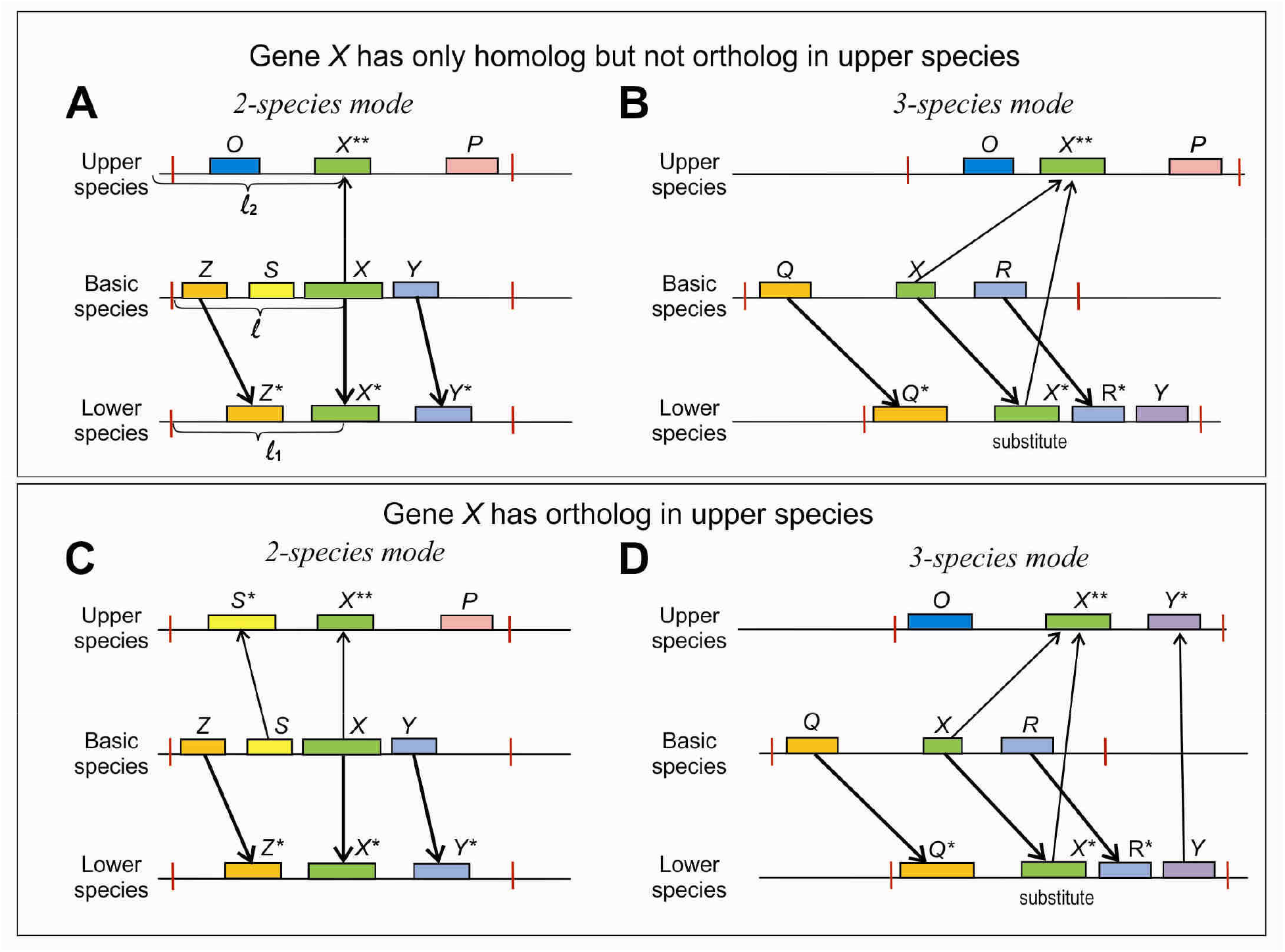
2- and 3-species modes of selection of genes having no orthologs in the upper species. **A and B**. The schemes demonstrate the case when gene *X* was considered as the lost one for 2- and 3- species modes respectively. Gene *X* has only homolog *X\*\**, but no ortholog in upper species, as the neighbors of *X* in basic (A) or lower species (B), *Z, S, Y*or *Q*, respectively, have no homologs in upper species, where gene *X\*\** neighbors are *O* and *P*. The red ticks indicate borders of the “window” in which local synteny was checked. The size of the window, l, was chosen in this particular work as2Mbp for all genes: *l=l_1_=l_2_*. Thin and bold arrows indicate homologous genes in the upper and lower species respectively. **C and D**. The schemes demonstrate the case when gene *X* was considered as preserved, for 2- and 3- species modes respectively. **Gene** *X* has ortholog *X\*\** in upper species, as at least one of the neighbors of *X* in basic (C) or lower species (D), *Z, S, Y* or *Q*, respectively, has homolog in upper species, where gene *X\*\** neighbors with *Y\** and *S\**.

In the 3-species mode, the following two conditions are regarded to state that gene *X* in the basic species has no ortholog in upper species (1) *X* has no witnesses near any of its homologs in upper species, (2) any ortholog of *X* in the lower species also has no witnesses near any of its own homologs upper species (Figure 1B). The computer implementation of this method is freely available at http://lab6.iitp.ru/ru/lossgainrsl. The program is deeply parallelized and can operate on a supercomputer, which is essential if great number of complete genomes are considered jointly or synteny blocks consist of many neighboring genes.

Let us denote the set of genes of the basic species obtained in the 2-species mode by the 2-species list and the one obtained in the 3-species mode by the 3-species list. The intersection of the 2- and 3-species lists is referred to as the gene list for given definitions of homology (see Materials and Methods). Thus, the intersection of the gene lists for specific definitions is referred to as the list of lost genes. It is the output of our method and program. Figures 1C and 1D illustrate the cases when the gene *X* in the basic species has orthologs in the upper species and thus, should be excluded from the final list of the lost genes.

For the data specified in the Supplementary materials section, our program predicted the following genes to be lost in warm-blooded animals: ENSXETG00000033176 (named *c-Answer* in this work), ENSXETG00000016048 (foxo1, forkhead box O1), ENSXETG00000006008 (E3 ligase, Prothymosin alfa related), ENSXETG00000023966 (*sfrpx*, secreted frizzled-related protein), ENSXETG00000025525 (*pnhd*, pinhead, secreted inhibitor of Wnt/bCatenin signaling), ENSXETG00000030282 (nuclear factor 7, zinc-finger protein), ENSXETG00000031627 (F-box protein), ENSXETG00000033120 (AP endonuclease 1), ENSXETG00000033543.

To confidently confirm that all of the detected genes have no orthologs in warm-blooded animals, they have all been checked for the absence of possible local synteny with genes in warm-blooded animals using the 5Mb window *l=l_1_=l_2_*.

Noteworthy, three genes missing in all placental mammals and in the majority of birds and reptiles (*Ag1*, *Ras-dva1* and *Ras-dva2*), which were empirically found by us earlier (Ivanova et al., 2013a, 2015; Tereshina et al., 2014), were also detected by our program as lost genes, if some species of warm-blooded animals, in which these genes are present, were excluded before processing. Thus, the results of this testing confirm the validity of our approach.

### Identified genes are activated during regeneration of the *X. laevis* tadpole hindlimb bud and tail

Temporal expression patterns of eight genes identified during the bioinformatic screening were analyzed in regeneration of *X. laevis* tadpole tail by qRT-PCR. In the experiment, we used tissue samples cut from stumps of tails on 0, 1, 2 and 6 days post amputation (dpa) where 0 dpa sample was the piece of stump harvested just after amputation. These 0 dpa samples were used as control, having the level of gene expression characteristic of the non-amputated appendages (Figure 2A). As a result, we detected significant increase of the expression level of four genes relative to the expression of two houskeeping genes, *EFalfa* and *ODC*, on 1 dpa in tail regenerates in comparison to 0 dpa (Figure 2A). By 6 dpa the expression levels of all genes returned to the respective basal levels. The revealed increase in expression of these four genes suggests their potential role in regeneration.

**Figure 2.**
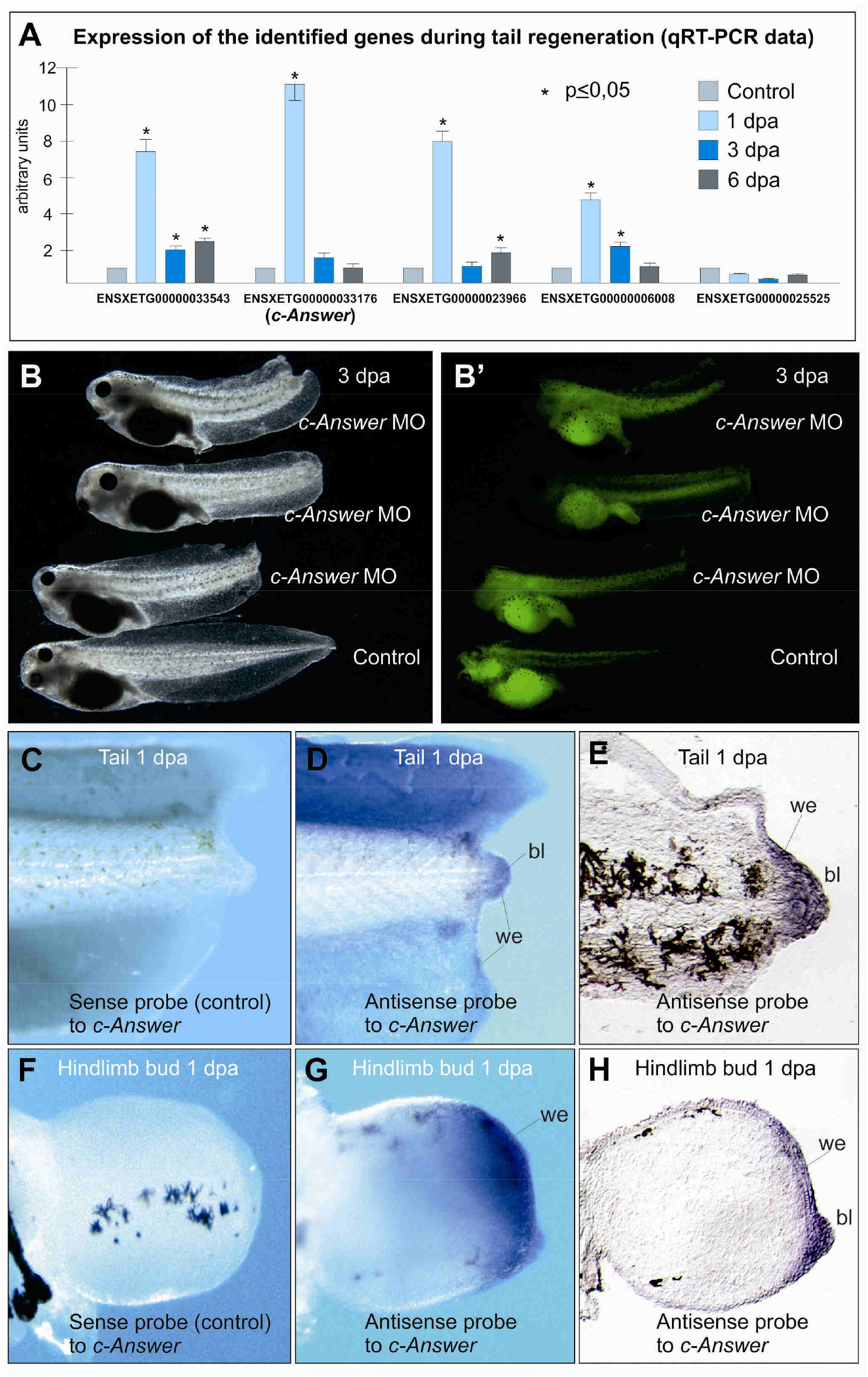
Analysis of expression and function of genes missing in warm-blooded animals in *X. laevis* tadpole tail regeneration. **A**. qRT-PCR analysis of *c-Answer* expression in blastema during tail regeneration at the indicated date after amputation. All results were normalized in relation to the geometric mean of two house-keeping genes, EF-alfa1 and ODC, expression as described (Ivanova et al., 2013). **B**. Injection of anti-sense morpholino oligonucleotides to *c-Answer* mRNA inhibits regeneration of the tadpole tail. **B.** Fluorescent image of same embryos as are shown on B demonstrates distribution of the co-injected tracer FLD. **C and F**. Whole-mount in situ hybridization with sense probe to *c-Answer* (control) of the amputated tail and hindlimb bud respectively at 1 day post amputation. **D and G**. Whole-mount in situ hybridization with anti-sense probe to *c-Answer* of the amputated tail and hindlimb bud respectively on 1 day post amputation. An increased expression of c-Answer is observed in the wound epithelium (we) and blastema (bl). **E and H**. Frozen histological sections of the regenerating tail and hindlimb bud respectively hybridized whole-mount with anti-sense probe to *c-Answer*.

Then, to verify if the identified genes are indeed critical for regeneration, we decided to test the effects of their down-regulation on the tadpole tail regeneration. To this end, we injected early *X. laevis* embryos with anti-sense morpholino oligonucleotides (MO) to mRNA of these genes (see results of testing the MO efficiency and specificity on Figure S1). The tadpole tails injected with MO were amputated at stage 41 and their regeneration capacity was compared. However, we were able to perform these experiments only in case of ENSXETG00000033176, ENSXETG00000033543 and ENSXETG00000023966. Injection of MOs for the rest five genes had dramatic effects that did not allow embryos to develop after gastrulation. Therefore, we concluded that the loss of these five genes in evolution could not be an initial cause of the regeneration capacity decrease due to the lethality of such events. Probably, these genes could be lost only at the next steps of the evolution, when the genetic mechanisms had already been significantly modified as a result of accumulation of some non-lethal mutations or loss of some other genes.

Among three genes, whose knockdown had no lethal effects, the most powerful retardation effect on tail regeneration was caused by ENSXETG00000033176 (Figures 2B and 2B’) . Due to its specific expression in the wound epithelium of the regenerating tail and hind limb bud (Figures 2C-H) and the ability of the protein to bind some membrane receptors (see below) we named this gene *c-Answer* (after <u>c</u>old-blooded <u>An</u>imals <u>s</u>pecific <u>w</u>ound <u>e</u>pithelial receptor-binding protein). Based on these results, we have chosen *c-Answer* for further investigation. ee results of testing the MO efficiency and specificity on Figure S1.

### c-Answer is a homodimer-forming transmembrane protein, homologous to FGF receptors

We have shown that *c-Answer* encodes a transmembrane protein, which has overall structure resembling that of the single-path receptors. In all tested species, *X. laevis* (GeneBank accession No; MG865735), *X. tropicalis* (MG865736), *Danio rerio* (MG865737), *Ambystoma mexicanum* (MG865738), c-Answer contains the amino terminal signal peptide, two Ig-like domains, single transmembrane helix and short cytoplasmic part (Figure 3A). While putative extracellular part of c-Answer has no high homology with any known proteins, it appears to be most homologous to Ig-like D2-D3 domains of the extracellular parts of receptors FGFR1-4, demonstrating highest homology with FGFR4 receptor (Figure 3A). Interestingly, these domains of FGFRs are critical for their binding to FGFs (Johnson et al., 1990; Lemmon and Schlessinger, 2010).

**Figure 3.**
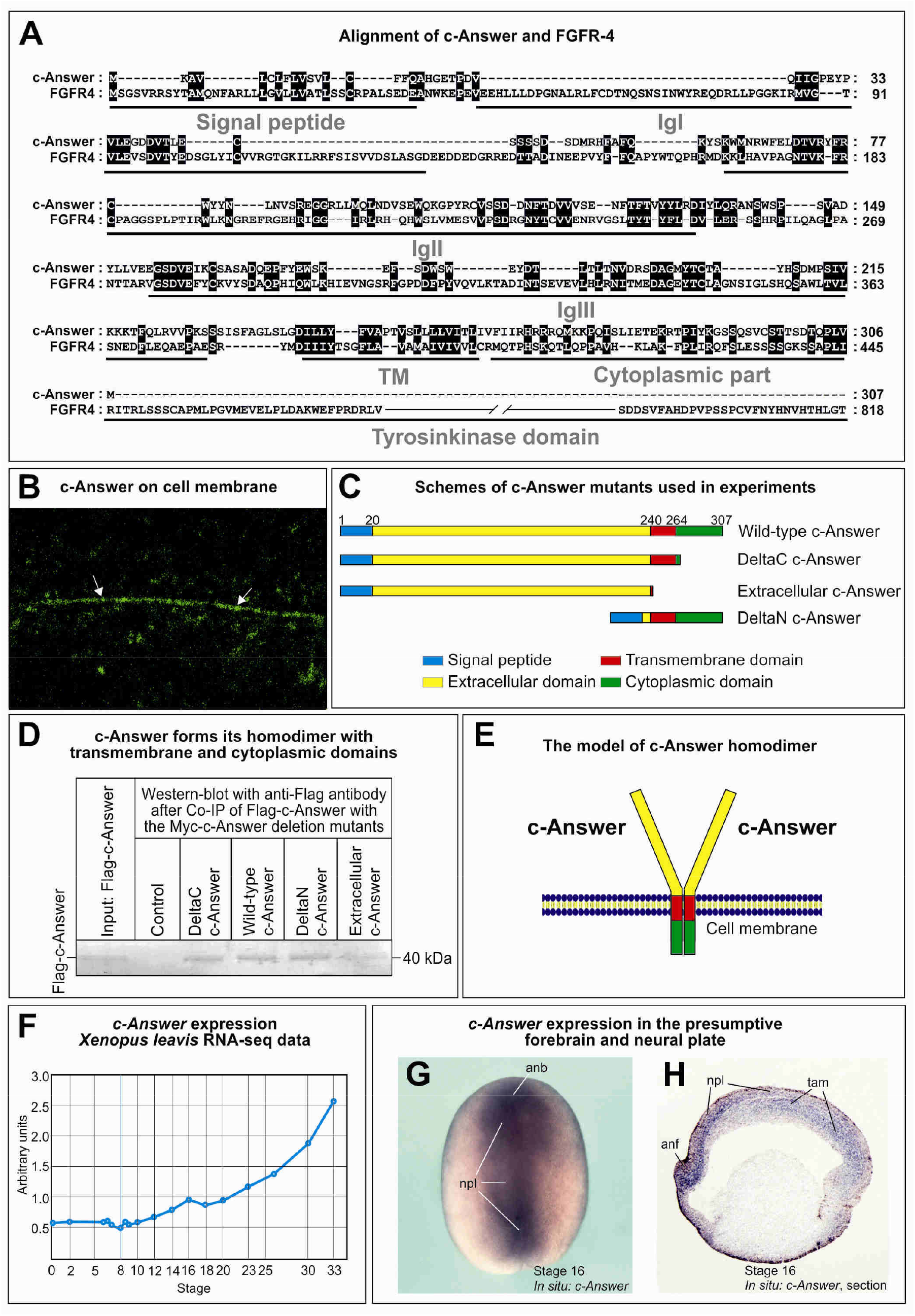
c-Answer encodes a transmembrane protein homologous to FGFR4, forming homodimer and ubiquitously expressed in the early embryonic development of *X. laevis*. **A**. Alignment of c-Answer with FGFR4. **B**. Localization of the secreted hybrid of EGFP and c-Answer on the membrane of the animal cap cell as revealed by confocal microscopy. **C**. Schemes of c-Answer deletion mutants used in experiments on co-immunoprecipitation and functional analysis. **D**. Western blotting with Flag-antibody after Co-IP of Flag-c-Answer with different Myc-tagged deletion mutants of c-Answer shown on C. **E**. The model of c-Answer homodimer according to the results of Co-IP shown on D. **F**. Temporal expression of c-Answer as revealed by RNA-seq analysis. **G and H**. In situ hybridization with c-Answer probe in whole-mounted middle neurula and on the frozen sagittal section of the embryo at the middle neurula stage.

Meanwhile, cytoplasmic part of c-Answer has no tyrosinkinase domains characteristic to FGFRs, although it contains one potential tyrosine phosphorylation site. Thus, c-Answer can be putatively considered a receptor lacking intrinsic kinase activity.

To confirm experimentally that c-Answer is a transmembrane protein, we investigated subcellular localization of its hybrid with EGFP. To this end, EGFP-Answer was translated from the synthetic mRNA injected into *X. laevis* embryos and, at early gastrula stage, its localization in the animal hemisphere was observed via confocal microscope. A significant portion of EGFP-c-Answer was localized to cell membranes (Figure 3B), confirming that c-Answer is a transmembrane protein.

It is well known that transmembrane proteins often form homodimers. Assuming this, we tested the ability of c-Answer to form a homodimer. The formation of the homodimer was detected by Co-IP in embryos co-injected with *c-Answer* mRNA tagged by Myc- and Flag-epitopes (Figures 3C and 3D). To understand which part of c-Answer is involved in the dimer formation, we investigated the ability of the following Flag-tagged deletion mutants of c-Answer to interact with the Myc-tagged wild-type c-Answer in Co-IP-test: extracellular c-Answer (c-Answer lacking transmembrane and cytoplasmic domains), deltaN-c-Answer (c-Answer lacking extracellular domain), deltaC-c-Answer (c-Answer lacking cytoplasmic domain) (Figure 3C). As a result, we established that primarily transmembrane and cytoplasmic domains of c-Answer are involved in the homodimer formation (Figures 3D and 3E).

### c-Answer is highly expressed in neurulation and appendage regeneration

According to a genome-scale database of mRNA dynamics in *X. laevis* embryogenesis, *c-Answer* transcripts are present at low level in the embryo till late gastrula stage, then their concentration begins to increase gradually (Figure 3F). Consistently with this, *c-Answer* transcripts were revealed by the whole-mount *in situ* hybridizations beginning from the late gastrula stage throughout the dorsal ectoderm, with a maximum of expression located within the neurectoderm (Figure 3G). As was shown by sectioning gastrula and neurula stage embryos, the expression of *c-Answer* is localized primarily in cells of the inner layer of the anterior neurectoderm and in the trunk axial mesoderm (**Figure 3H**).

The results of single-cell RNA sequencing obtained by (Briggs et al., 2018) also support our experimental data on *c-Answer* expression. Though, according to single-cell data, *c-Answer* expression at low level distributes rather uniformly and is present in all germ layers (ectoderm, mesoderm, and endoderm) some tissue subtype clusters, especially those of the presumptive forebrain and epithelial tissues are enriched in *c-Answer*. (Figure S5)

Consistently with the results of qRT-PCR analysis, an increase in *c-Answer* expression was seen during regeneration in stumps of both tails and hindlimb buds already at 1 dpa. Importantly, in both cases, *c-Answer* is expressed in the wound epithelium, which indicates possible involvement of *c-Answer* in regulation of this tissue critical for regeneration (**Figures 2D-H**).

In sum, the results of in situ hybridization, qRT-PCR experiments and single-cell sequencing data indicate that *c-Answer* could be involved in the early development of the forebrain, as well as in the body appendages regeneration.

### c-Answer down-regulation and overexpression have opposite effects on the brain development and regeneration

To investigate the physiological function of *c-Answer* during CNS development and tail regeneration, we first analyzed effects of its down-regulation by injection of anti-sense morpholino oligonucleotides to *c-Answer* mRNA and by CRISPR/Cas9 knockout of *c-Answer* gene (see results of the embryo genotyping after CRISPR/Cas9 procedure in **Figure S2**). As a result, in both cases diminishing of the overall tadpoles size was observed (**Figures 4A and 4B**). Especially this concerned diminishing of the forebrain, including eyes and cement gland, and was most evident when MO or CRISPR/Cas9 were injected in only one of the two blastomeres at 2-cell stage (**Figures 4A and Figure S4B**). At the same time, a retardation of tail regeneration was observed in tadpoles, stage 41, injected with the MO or CRISPR/Cas9 (**Figures 4C-E**). Thus, we concluded that down-regulation of *c-Answer* leads to both the reduction of the forebrain size and retardation of tail regeneration. No such effects were observed in embryos injected with control, mis-c-Answer MO (not shown).

**Figure 4.**
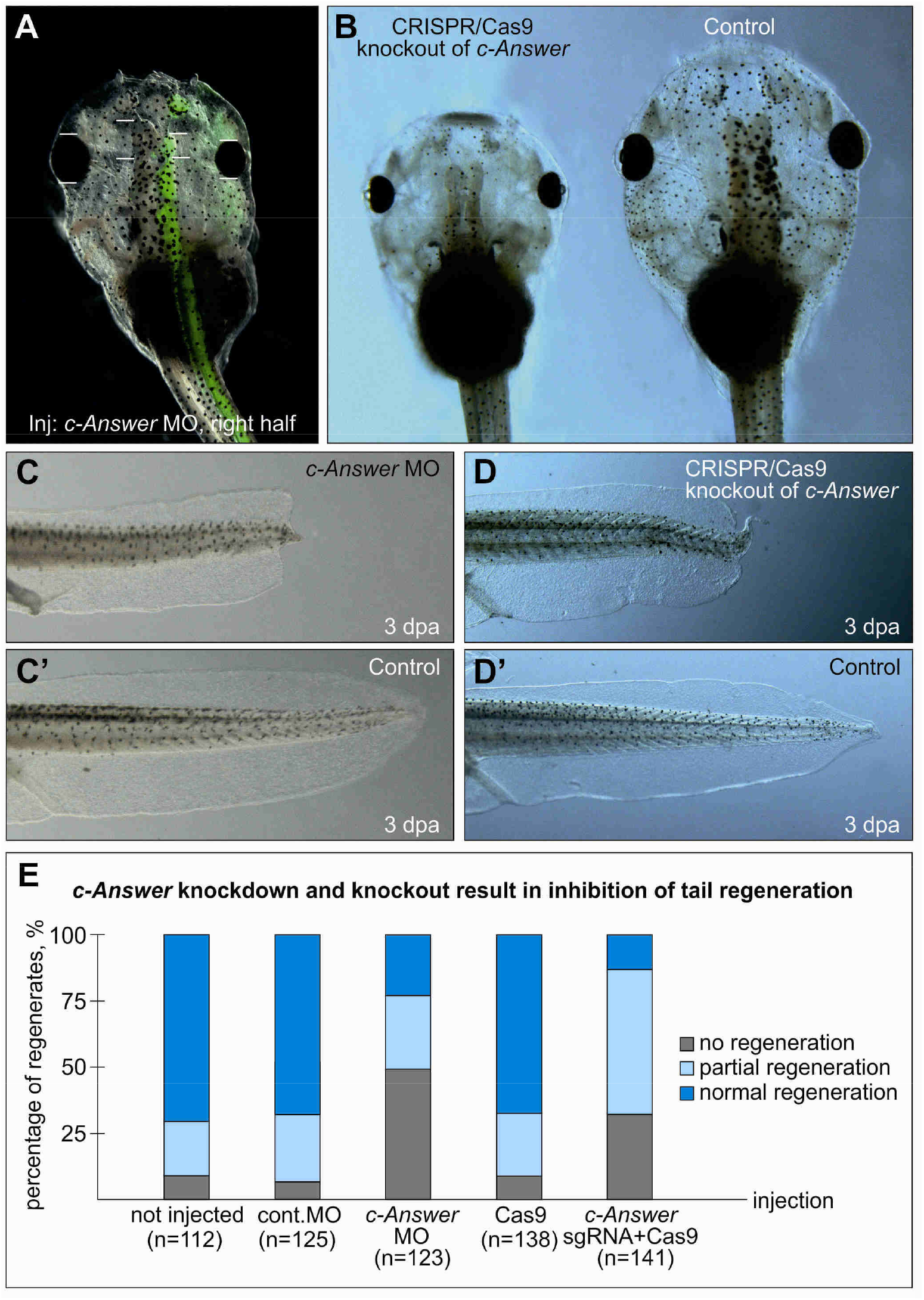
Effects of *c-Answer* downregulation by knockdown with anti-sense morpholino oligonucleotides and CRISPR/Cas9 knockout on the tadpole brain development and tail regeneration. **A**. *c-Answer* knockdown with *c-Answer* MO injections into the dorsal right blastomere at 4-cell stage results in the diminishing of the overall tadpole size, especially of the forebrain and eye as compared to the left side (control). Overlay with the fluorescent image demonstrates the distribution of the co-injected tracer, FLD. **B**. Tadpole in which c-Answer was knocked out with CRISPR/Cas9 technology has smaller size then the wild-type tadpole at the same stage. **C. C’ and D, D’**. Tail regeneration in tadpoles with the *c-Answer* knockdown or knockout, respectively, is inhibited in comparison to the wild-type control. **E**. Diagram showing the distribution of tail regeneration phenotypes in tadpoles injected with the indicated MO or components of CRISPR/Cas9 system.

Given the fact that *c-Answer* is expressed in the presumptive forebrain region at the very beginning of its specification, i.e. at the early neurula stage, we decided to verify if the observed forebrain malformations were caused by the lack of *c-Answer* activity during neurulation. To this end, we studied the effects of the *c-Answer* MO injections on expression of the forebrain specification genes at the midneurula stage. Consistently with the effects observed in tadpoles, a moderate reduction of the telencephalic regulator *FoxG1* and eye regulators *Rx1* and *Pax6* was detected (**Figures 5A-B and Figure S3A**). Interestingly, in all cases, the areas in which down-regulation of *c-Answer* caused inhibition of regulator-gene expression were located in the lateral regions of their normal expression domains, corresponding to the presumptive dorsal part of the telencephalon and eyes. No inhibitory effects were seen in the medial regions, which give rise predominantly to the ventral parts of the telencephalon and eyes. No effects of *c-Answer* knockdown were revealed in case of *Six3* that is expressed throughout the anterior neural ectoderm, and *En1*, which expression marks the presumptive mid-hindbrain border (Figures S3E and S3H). These results confirm specificity of the *c-Answer* function.

**Figure 5.**
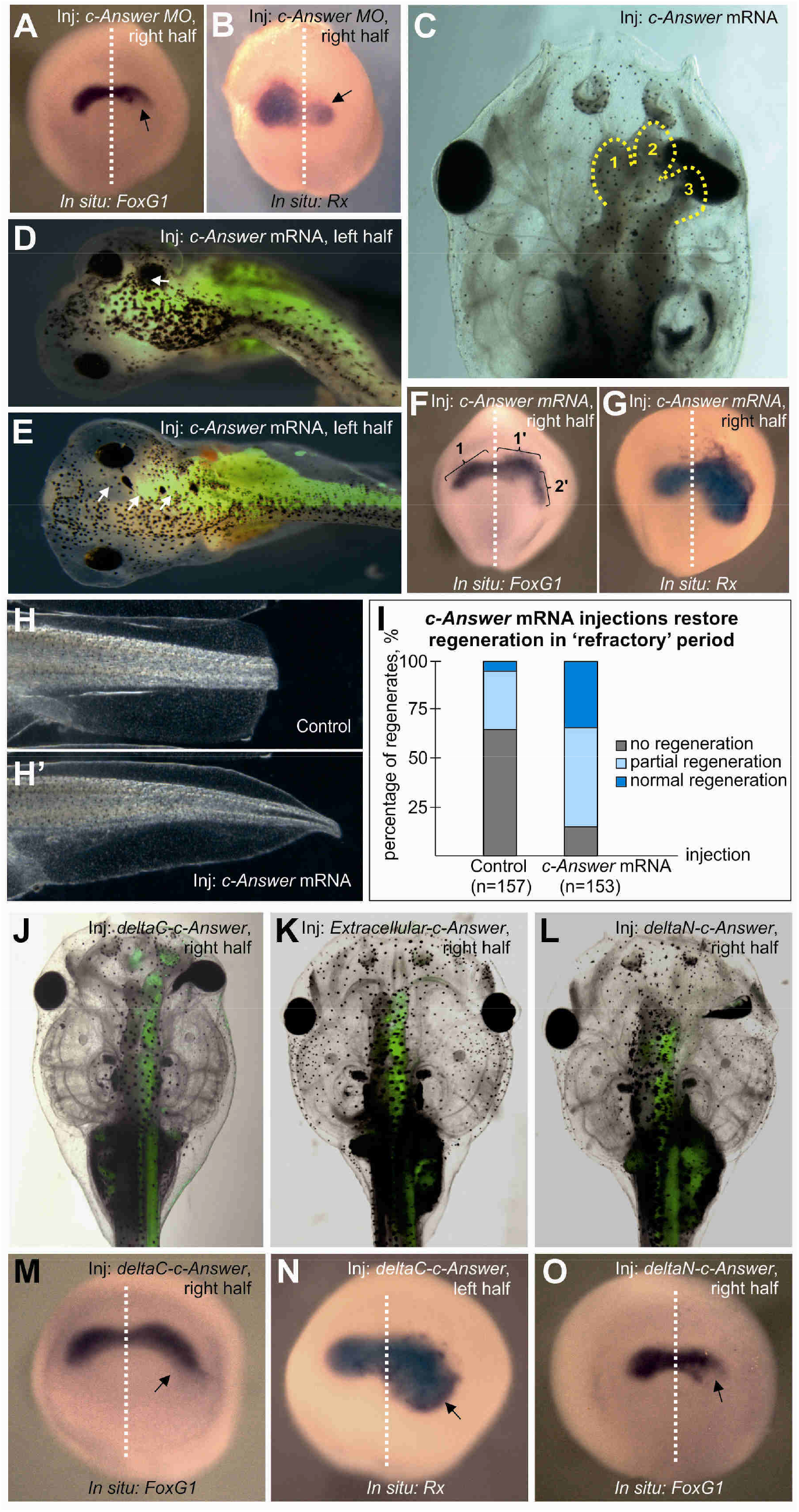
Effects of c-Answer and its deletion mutants overexpression on the tadpole brain development and tail regeneration. **A and B**. Inhibition of *FoxG1* and *Rx* expression on the right side of the middle neurula embryos injected into the right dorsal blastomere at 4-cell stage. **C**. Overexpression of wild-type *c-Answer* results in the development of ectopic telencephalic hemisphere (3) in addition to normal ones (1 and 2). **D and E**. Ectopic eye differentiation in tadpoles overexpressing wild-type *c-Answer*. **F and G**. Ectopic expression of *FoxG1* and *Rx* on the right side of the middle neurula embryos injected into the right dorsal blastomere at 4-cells stage with *c-Answer* mRNA. **H and H’**. Overexpression of the wild-type *c-Answer* rescues the tail regeneration in the ‘refractory’ period. **I**. Diagram showing the distribution of the regenerating tail phenotypes in the control tadpoles and those overexpressing wild-type *c-Answer*. Tails were amputated in the ‘refractory’ period. **J**. Overexpression of deltaC-c-Answer mutant leads to the telencephalon size increase and ectopic RPE differentiation. **K**. Overexpression of the extracellular domain of c-Answer resulted in a slight increase in the telencephalic and eye size on the injected (right) side. **L**. Overexpression of deltaN-c-Answer mutant inhibits development of the telencephalon and eye on the injected side. **M and N**. Ectopic expression of *FoxG1* and *Rx* on the right side of the middle neurula embryos injected into the right dorsal blastomere at 4-cell stage with deltaC-c-Answer mRNA. **O**. Inhibition of *FoxG1* expression in the lateral part of the endogenous right expression domain of *FoxG1*.

Then, we investigated *c-Answer* gain-of-function effects by the unilateral overexpression of *c-Answer* mRNA. As a result, two types of effects were revealed in the head region. First, an increase of the telencephalic differentiation, ranging from slight increase of the telencephalon on the injected side to the additional part of the telencephalon, was detected in 70% of embryos (52/74) (**Figure 5C, Figures S4C and S4C’**). In addition, ectopic cement glands were frequently observed in these embryos (**Figures S4C**). Second, a range of eye phenotypes was observed on the injected side in almost all these embryos (**Figure 5D and 5E, Figure S4C-G**). These phenotypes included disrupted eye development; ectopic retinal pigment epithelium (RPE); RPE extensions (**Figure 5J and 5L, Figure S4D-F**); expansion of RPE and positioning of eye cup adjacent to forebrain and even development of secondary eye in rare cases (**Figure 5D, Figure S4G**). Importantly, these malformations were accompanied by the expansion of the expression zones of the telencephalic regulator *FoxG1* and eye regulators *Rx1* and *Pax6* and during neurulation (**Figures 5F and 5G, Figures 2B and 2C**). However, as in case of *c-Answer* knockdown, no effects were detected for *Six3* and *En1* (**Figures S5F and S5I**).

To verify that *c-Answer* overexpression could influence tail regeneration, we studied the ability of exogenous *c-Answer* to rescue the regeneration during the so-called ‘refractory’ period (stages 45-47), when the regeneration of tail appears to be blocked due to natural reasons using the previously developed approach (Ivanova et. al. in press). As a result, we observed a significant increase in the number of tadpoles with the restored tail regeneration capacity during the ‘refractory’ period (**Figures 5H-I**’). Obviously, this indicates that *c-Answer* can stimulate regeneration in the ‘refractory’ period.

Noteworthy, the results obtained in *X. laevis* embryos with *c-Answer* knock-down and knock-out demonstrating the forebrain, eyes and cement gland development inhibition are consistent with the single-cell sequencing data, according to which *c-Answer* is mostly represented in cement gland primordium, anterior neural tube, adenohypophyseal placode, and eye primordium at stages 16, 18 and 22 (**Figure S5**).

### c-Answer deletion mutants require transmembrane domain to influence CNS development and regeneration

As c-Answer has extracellular and cytoplasmic parts, we decided to test what effects may be caused by its deletion mutants lacking various domains, in order to shed light on c-Answer functioning at molecular level. Given this, we investigated the effects of Extracellular c-Answer (c-Answer lacking transmembrane and cytoplasmic domains), deltaN-c-Answer (c-Answer lacking extracellular domain), deltaC-c-Answer (c-Answer lacking cytoplasmic domain) mutants on the brain development and tail regeneration (**Figure 3C**).

When deltaC mRNA was injected, we observed the effects resembling those of the wild-type c-Answer overexpression. Namely, the increase of the telencephalon in embryos unilaterally injected with deltaC mRNA and ectopic differentiation of eye pigment epithelium (65%, 54/83) (**Figure 5J**). However, in contrast to wild-type *c-Answer* mRNA injections, differentiation of ectopic pigment epithelium was always observed in vicinity of the normal eye, particularly, around the optic nerve. Ectopic retinal pigment epithelium differentiation was never detected far from the normal eye, like it was in case of the wild-type c-Answer mRNA injections. Also, well-structured secondary eyes were never detected in these embryos. At the same time, the eye on the side injected with deltaC mRNA was frequently increased, as in embryos injected with the wild-type *c-Answer* mRNA (**Figure 5J**). Consistently to the increased telencephalon and eyes, an expansion of *FoxG1, Pax6* and *Rx* expression was detected on the injected side at neurula stage (**Figures 5M and 5N, Figure S5D**). Thus, deltaC mutant injections resembled the wild type *c-Answer* mRNA injections, but differed from the latter by more severe and frequent increase of the telencephalon and less ectopic eye differentiation.

The effects of deltaC-c-Answer upon tail regeneration resembled those of *c-Answer* MO. Namely, a retardation of regeneration was seen in tadpoles developing with *deltaC-c-Answer* mRNA injected into blastomeres that give rise to tail (28% no regeneration, 54% partial regeneration. 18% normal regeneration; 36/70/24).

Given that deltaC mutant of c-Answer, containing its extracellular part and the transmembrane domain, caused effects resembling those of the wild-type c-Answer, we thought to test if the same effects could be elicited by the extracellular part of c-Answer (Extracellular mutant). However, no brain outgrowth and ectopic eye differentiation, as in case of wild-type Answer or its deltaC mutant, was observed. At the same time, slight increase in the telencephalon and eye size was seen in 30% of the injected tadpoles (25/83) (**Figure 5K**). Additionally, we have not observed any retardation in tail regeneration.

Therefore, we conclude that while the extracellular part of c-Answer is necessary for the interaction with P2Y1 and FGFR4 receptors, it cannot effectively operate without the transmembrane domain.

Then we investigated effects of c-Answer mutant lacking the extracellular part (deltaN mutant). As a result, a set of abnormalities resembling those elicited by *c-Answer* MO, including the telencephalon and eye size diminishing (75%, 60/80) and retardation of regeneration (24% no regeneration, 49% partial regeneration, 27% normal regeneration; 29/59/32), were observed (**Figures 5L and 5O**). Obviously, this indicates that deltaN mutant of c-Answer is likely to operate as its dominant-negative mutant.

In summary, the data obtained indicate that transmembrane part of c-Answer is essential for its activity. Isolated Extracellular domain has no significant effect on brain development and regeneration. In turn, c-Answer deprived of extracellular (deltaN) or cytoplasmic part (deltaC) operates as dominant-negative or dominant-positive mutant, respectively, that can seemingly compete with the wild-type c-Answer.

### c-Answer interacts with FGFR1-4 and P2Y1, but not with Fgf8

Assuming the results of experiments on c-Answer misexpression, as well as its potential function as the transmembrane protein, one may suppose that it could be involved in signaling that regulates early stages of the telencepalic and eye differentiation. We focused on two types of such signaling, as they both operate in the anterior neural plate and their disruption has effects similar to those observed in c-Answer misexpression experiments. The first one is Fgf8 signaling that activates expression of the telencephalic master regulator *FoxG1* in cells of the Anterior Neural Border (ANB) (Danesin and Houart, 2012a; Houart et al., 1998; Shimamura and Rubenstein, 1997). The second is ADP purinergic signaling through transmembrane receptor P2Y1. Like overexpression of wild-type *c-Answer*, overexpression of *Fgf8* expands expression zone of *FoxG1* and increases the telencephalon size, while overexpression of *P2Y1* induces ectopic eye differentiation as it was observed in case of wild-type *c-Answer* mRNA injections (Massé et al., 2007).

Given this correlation of misexpression effects, we decided to test whether c-Answer can directly interact with extracellular protein components of the Fgf8 and purinergic signaling pathways. To this end, we arranged experiments on co-immunoprecipitation of Myc epitop-tagged c-Answer with Fgf 8a/b, the receptors of Fgf8, FGFR1-4, and the receptor of ADP, P2Y1, tagged by Flag epitop. All these proteins were translated from the corresponding synthetic mRNA injected in pairs into the early *X. laevis* embryos (see Materials and Methods). As a result, we established that in these conditions c-Answer binds to all FGF receptors and P2Y1, but not to Fgf8 (Figures 6A and 6B).

**Figure 6.**
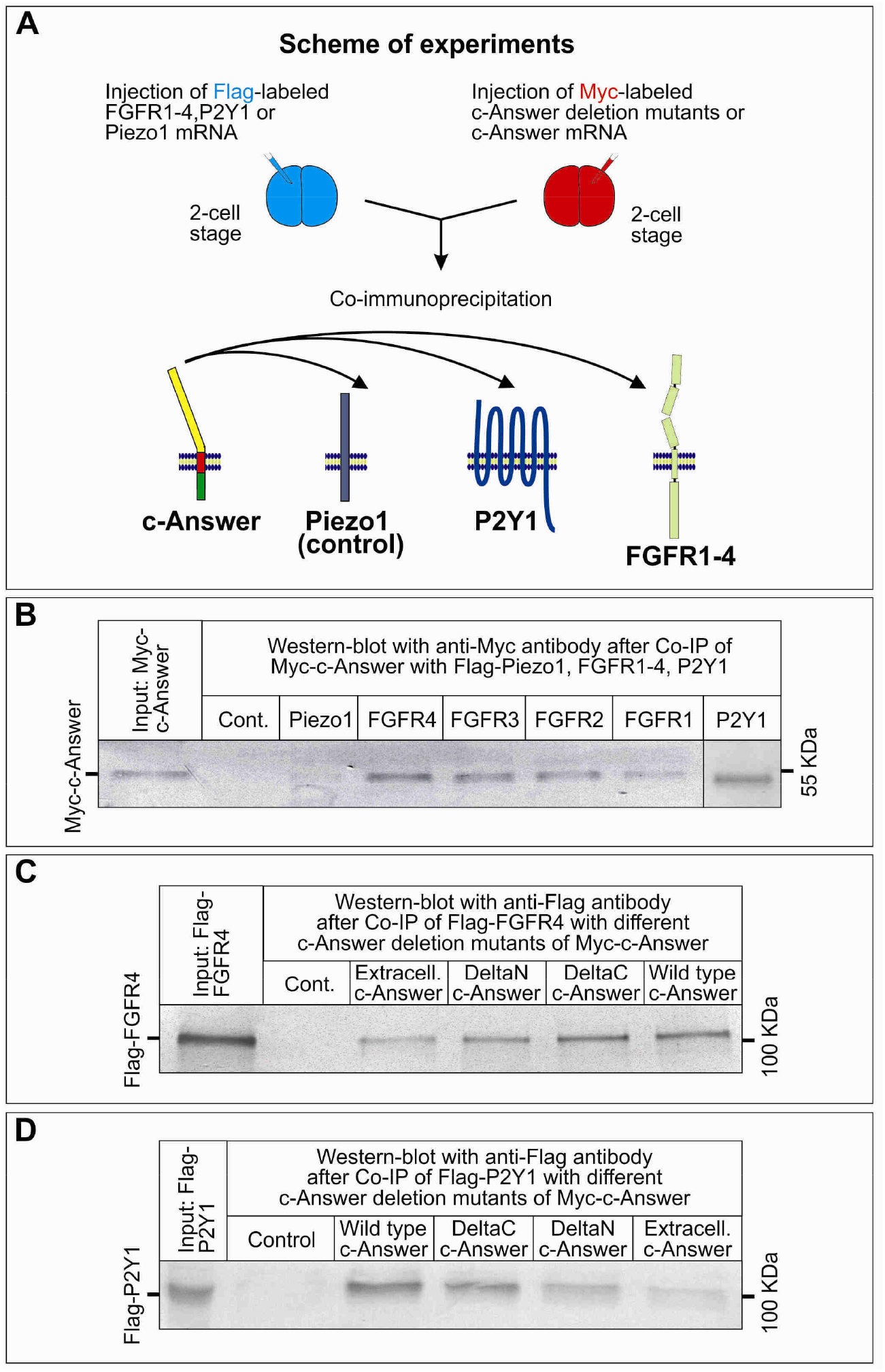
Analysis of c-Answer and its deletion mutants binding capacity with FGFR1-4 and P2Y1 receptors. **A**. Scheme of experiments. Flag- and Myc-tagged proteins were separately expressed from synthetic mRNA in embryos, embryonic extracts were mixed for Co-IP and analyzed by Western blotting. **B-D**. Western blotting analysis with anti-Myc or anti-Flag antibodies after the indicated Co-IP.

Then, to understand which domains of c-Answer are responsible for its interaction with FGFR4 and P2Y1, we investigated the ability of Myc-tagged deletion mutants of c-Answer to bind Flag-tagged receptors in Co-IP test. As a result, we revealed that all of the tested deletion mutants could interact with FGFR4 and P2Y1 albeit with a weaker affinity than wild-type c-Answer (**Figures 6C and 6D**). At the same time, somewhat stronger interaction was observed with deltaC and deltaN mutants which contained the transmembrane domain. Thus, we concluded that all domains of c-Answer are to some extent involved in its interaction with the receptors.

### c-Answer promotes FGF and P2Y1 signaling

To understand how c-Answer can influence FGF signaling, we tested effects of c-Answer upon downstream signaling pathways activated by Fgf8. To this end we used pGL4.33 (SRE) reporter vector (Promega) in which the fire-fly luciferase was driven by response elements sensitive to MAP/ERK pathway, i.e. the key pathway activated through tyrosinkinase receptors, including FGF receptors. Another reporter, pGL4.44 (AP1) (Promega), sensitive to the stress-activated MAPK/JNK pathway, that cannot be activated by growth factors, was used as a control.

In these experiments, we injected embryos with pGL4.33 (SRE) reporter in a mixture with either *Fgf8* mRNA, or *Fgf8* and *c-Answer* mRNAs, then cut animal caps off the injected embryos at early gastrula stage and analyzed the luciferase signal at late gastrula stage equivalent

(**Figure 7A**). As a result, we observed an increase of MAP/ERK in cells co-injected with *Fgf8* and *c-Answer* mRNA (Figure 7B). In contrast, no activation of MAPK/JNK pathway was seen in similarly arranged control experiments (**Figure 7B**). Importantly, lower signal of the reporter was detected in case when c-Answer was expressed alone or in the animal caps injected only with the reporter (**Figure 7B**). Thus, we concluded that c-Answer promotes MAP/ERK pathway activated by Fgf8 signaling.

**Figure 7.**
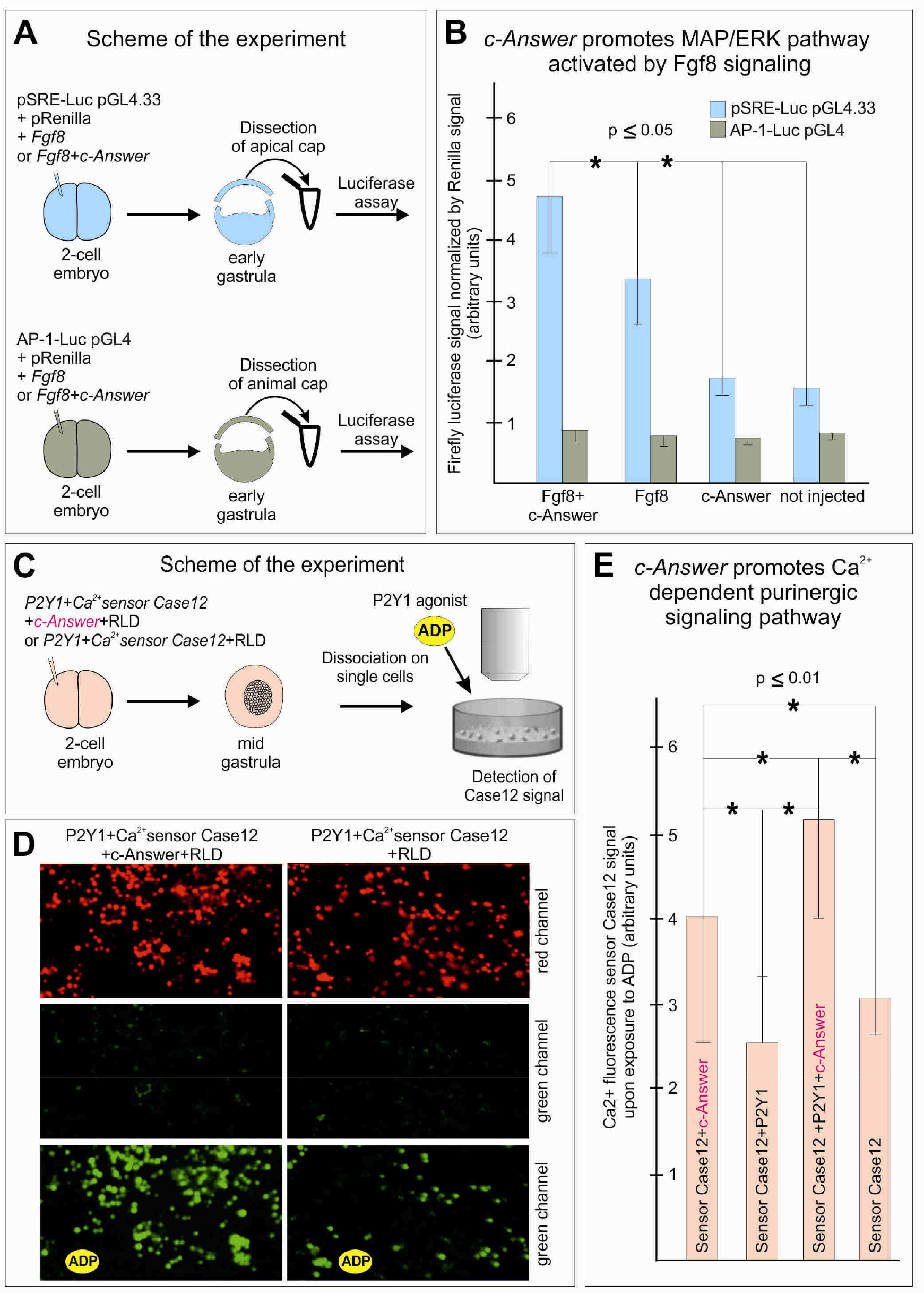
c-Answer promotes signaling through FGFR4 and P2Y1 receptors. **A**. Scheme of the experiment on the analysis of c-Answer effects upon expression of MAP/ERK pathway luciferase reporter pSPE-Luc pGL4.33. Stress-activated reporter AP-1-Luc pGL4 was used as a control. **B**. Diagram showing the results of Luc signal analysis for two reporters in animal caps of embryos expressing the indicated proteins. **C**. Scheme of experiment on the analysis of c-Answer effects upon Ca^2^+ flux in response to addition of P2Y1 agonist, ADP, to the animal cap cells expressing Ca^2^+ sensor Case12 and purinergic receptor P2Y1. **D**. Fluorescent images of cells expressing the indicated proteins before and after ADP addition. **E**. Diagram showing the results of Case12 signal analysis in animal cap cells expressing the indicated proteins.

Given that a key cytoplasmic mediator of purinergic signaling is Ca^2+^, we arranged experiments with Ca^2+^ fluorescent reporter to test effects of c-Answer on this signaling through P2Y1 receptor. To this end, we injected embryos with mRNA encoding for Ca^2+^-sensitive variant of green fluorescent protein (Case9 reporter protein, Evrogen) mixed with either *P2Y1* mRNA, or with *P2Y1* and *c-Answer* mRNAs. Animal caps of these embryos were dissociated into single cells in Ca^2^+-free medium and then Ca^2^+ flux into cytoplasm of these cells after application of ADP was monitored via measuring the fluorescence of Case9 reporter (**Figure 7C**). As a result, we revealed higher fluorescent signal in cells expressing exogenous c-Answer (**Figure 7C, Video S1**). Similar results were obtained in whole embryos (**Video S2**).

## DISCUSSION

### A wide-range bioinformatic screening for genes lost in higher vertebrates introduces novel way of orthologous genes identification

A widely accepted point of view is that the rearrangement of gene cis-regulatory elements network is the main type of the evolutionary changes that led to the phenotypic and physiological differences between species, in particular in vertebrates (Rodríguez-Trelles et al., 2003; Wray, 2007). Unfortunately, this kind of genomic rearrangements are difficult to reveal in frames of systematic screening by bioinformatics methods, because of uncertainty during identifying cis-regulatory elements and revealing of how mutations in their content are connected with specific changes in gene functioning.

Meanwhile, there is another type of mutations, where relation to specific phenotypic or physiological changes could be more easily revealed, namely, mutations within gene sequences coding for proteins. Among them, loss-of-function mutations, which include nonsense mutations or insertion/deletion mutations that lead to the frame shift followed by the production of nonfunctional proteins, are of special interest because they may lead to complete loss of particular genes. Obviously, the majority of loss-of-function mutations should be lethal. However, some of them may still provide a benefit for the organism. In that case, these mutations will be fixed by the natural selection, which in turn should lead to a quick loss of the rest of the gene sequences due to acceleration of the mutation process in the corresponding genomic regions under these conditions. Importantly, although such complete physical loss of gene should be extremely rare event due to its harmful effect, it can be easily revealed by bioinformatic methods as compared to any other type of mutation.

The rationale for search for such genes is based on the assumption that phenotypic or physiological difference between different classes of animals in some way might be caused by the loss of certain important genes. This is confirmed by our recent empirical finding that genes, Ag1, *Ras-dva1* and *Ras-dva2*, were lost in poorly regenerating higher vertebrates, but still play important roles during the body appendages restoration in well regenerating fishes and amphibians (Ivanova et al., 2013a, 2015; Tereshina et al., 2014).

In the present work, we have developed an algorithm and a computer program for systematic search for genes that were lost at a certain step of the evolution. During this screening, we considered the lost genes to be those that have no direct homologs (orthologs) in all the species located on the evolutionary tree above the selected species, but do have such orthologs in species below it. Two criteria were used to identify orthologous genes. First, high homology of proteins encoded by the analyzed genes and second, the demand of the local genomic synteny. The introduced requirement for local genomic synteny is necessary to ensure that the two genes from different species are indeed orthologs, as it is the only requirement that may guaranty the presence of the same set of cis-regulatory elements in both genes with high probability. Alternatively, even in case of high homology between proteins, the coding region of the non-orthologous gene will be surrounded by different non-coding sequences, which contain different set of cis-regulatory elements. Thus, despite high homology of its protein, this non-orthologous gene in one species will be unable to completely substitute functions of its homolog in other species, and this is just why it cannot be considered a true ortholog of the latter.

However, the developed algorithm obviously has certain technical limitations, which make it impossible to detect evolutionary gene loss in certain cases. Firstly, when highly homologous genes form a cluster, being located in a vicinity of each other in the same chromosome. In such case, loss of one of these genes at some evolutionary step cannot be detected by this algorithm, due to the presence of other homologs in the same locus and preservation of the local synteny for all of them. Secondly, for species with poorly sequenced genomes it appears impossible to carefully investigate local synteny for some genes and thus, such genes should be simply excluded from the analysis at all. One may be confident, however, that in all cases when well sequenced genomes are available, the genes selected by the developed algorithm as the lost at certain step of evolution, were indeed lost at this step.

As a result of the wide-range computer screening by the developed method for the genes lost in warm-blooded vertebrates, we were able to identify only eight genes. Even assuming the technical limitations of the method discussed above, the number of the lost genes seems to be surprisingly low. Obviously, this confirms that the complete gene deletion is quite rare phenomenon and phenotypic and physiological difference between different classes of vertebrates are mostly based on rearrangement of the genomic regulatory networks, but not on changes in gene repertoire.

Interestingly, at least four of the identified genes have demonstrated evident activation of expression at the very beginning of tadpole body appendage regeneration. However, only down-regulation of *c-Answer* affected regeneration specifically; down-regulation of three other genes caused severe overall damage to the embryo. This may indicate that the loss of these genes in evolution could not be an initial cause of the regeneration ability decrease due to the lethality of such events. Probably, these genes could be lost only at the next steps of the evolution, when the genetic mechanisms were already significantly modified as a result of accumulation of some non-lethal mutations or loss of some other genes.

### *c-Answer* is a novel regulator of the forebrain development and body appendage regeneration in *X. laevis*

*c-Answer* encodes a transmembrane protein, having the primary structure most homologous to FGFR4, especially to its D2 and D3 Ig-domains, transmembrane and juxtamembrane domains. However, in contrast to FGFRs, c-Answer has no D1 Ig-domain in the extracellular part as well as a tyrosine kinase domain in the cytoplasmic part. As we have shown, overexpression of *c-Answer* results in an increase in the telencephalic and eye differentiation, ranging from RPE extension to ectopic eye emergence in rare cases. These effects are accompanied by moderate increase of the expression zones of telencephalic regulator, *FoxG1*, and eye regulators, *Rx* and *Pax6*. In contrast, down-regulation of *c-Answer* elicits reduction of the telencephalon and eye size. Consisting with these results, that indicate involvement of *c-Answer* in regulation of telencephalon and eye development, we have demonstrated the ability of c-Answer to interact with FGFR1-4 and P2Y1 receptors, which transmit this signaling. Stimulating effects of c-Answer on the activity of these receptors was confirmed at physiological and molecular levels. At the same time, we were unable to reveal the interaction of c-Answer with the key inducer of the *FoxG1* expression and the telencephalic differentiation, Fgf8. This indicates that though c-Answer stimulates Fgf8 signaling via FGFR1-4, it cannot interact directly with Fgf8 at all, or at least is unable to interact with the latter in the absence of FGFR1-4. One may suppose that the same is also true in case of stimulating influence of c-Answer on eye differentiation trough modulation of P2Y1 signaling. However, this has to be specially tested in the future.

Using tadpole tail regeneration model, we determined that besides telencephalon development, *c-Answer* is necessary for appendage regeneration. Keeping in mind that Fgf signaling, including Fgf8 and FGFR4, is a key regulatory component of the regeneration molecular machinery (Gorsic et al., 2008; Lin and Slack, 2008), one may suppose that as well as in the telencephalon development, c-Answer plays a role of positive modulator of Fgf8 signaling during regeneration. We supposed c-Answer could play a role of an enhancer of Fgf8 signaling by compensating low level of Fgf8 and its receptor in the earliest period of regeneration. Furthermore, in conjunction with this, one may suppose that the loss of *c-Answer* in warm-blooded animals could be one of the factors that reduced their body appendages regenerative capacity.

In contrast to Fgf signaling, there is no direct evidence in the literature that purinergic signaling through P2Y1 is involved in regeneration. At the same time, crucial role of P2Y receptors was shown in the wound healing (Greig et al., 2003; Iwanaga et al., 2013). Therefore, one could not exclude that positive modulation of P2Y 1 by c-Answer revealed in the present work is important for regeneration.

Importantly, among a number of known transmembrane proteins that modulate activity of Fgf receptors, Sef and XFLRT3 were shown to operate like c-Answer during gastrulation and neurulation in *X. laevis* embryos and to interact with FGFRs inhibit or promote their activity respectively (Böttcher et al., 2004; Tsang et al., 2002). The authors speculate that Sef and XFLRT3 modulate FGFR signaling either directly or recruiting some cytoplasmic cofactors. Same modes of activity probably could be also valid for c-Answer interaction with FGFR1-4 and P2Y1. However, as c-Answer, in contrast to Sef and XFLRT3, binds to two different types of receptors, one may suppose that different regions of its molecule are responsible for binding to each of these receptor types.

### Possible role of c-Answer in the forebrain development and in evolution

As we established, c-Answer is expressed predominantly within the anterior neuroectoderm and positively regulates signaling through FGFR1-4 and P2Y1 receptors, which induce telencephalic and eye development respectively. Notably, though being comparatively low during gastrulation, the expression level of c-Answer progressively increases throughout the anterior neural plate by the end of neurulation. This indicates that stimulating effect of c-Answer on FGFR1-4 and P2Y1 signaling should be as well increasing progressively during neurulation. Given this, one may suppose that elevation of c-Answer concentration during neurulation is a signal that coordinates activation of the telencephalic and eye differentiation at specific period of development in different regions of the anterior neural plate, which is possible due to unique capability of c-Answer to interact with two different receptors. When the concentration of c-Answer is low, the level of FGFR1-4 and P2Y1 signaling is probably also lower some threshold necessary to start the telencephalic and eye differentiation. However, elevation of c-Answer concentration, which occurs simultaneously in the presumptive telencephalic and eye regions, could shift FGFR1-4 and P2Y1 signaling levels above the threshold thereby stimulating differentiation in both regions. We suggest, that coordination of telencephalic and eye differentiation stimulated by *c-Answer* may be necessary to stabilize the development. In turn, down-regulation of *c-Answer* may lead to misbalance of differentiation, which may result in a desynchronization between the telencephalic and eye development. Assuming that *c-Answer* was lost in the ancestors of warm-blooded animals, one may further hypothesize that heterochronies in development of different structures within the anterior neural plate, which might appear due to such desynchronization, could provide benefit for the progressive evolution of the warm-blooded animals forebrain, as a tradeoff for the reduced regeneration capability. In support of this, the work can be cited (Cavodeassi and Houart, 2012), where it was supposed that the early start of the telencephalic differentiation in mammals compared to fishes could result in relatively lager territory of telencephalic differentiation in mammals.

Another possible benefit that warm-blooded vertebrates might get as a result of *c-Answer* loss, could be a decline of the expression level of *FoxG1* in the dorsal region of the telencephalon. It was shown in mouse that a reduced level of *FoxG1* expression in the dorsal telencephalon is critical for the development of this region, which is predominantly increased during evolution of higher vertebrates (Danesin and Houart, 2012b). As we have found out, down-regulation of *c-Answer*, although not entirely eliminated *FoxG1* expression, elicited a significant decline of its expression level, especially in the presumptive dorsal regions of the *FoxG1* expression territory (**Figure 5A**). Thus, one may speculate that the loss of *c-Answer* in the ancestors of cold-blooded animals could result in a decline of *FoxG1* expression in the dorsal telencephalon, which in turn might provide conditions for the progressive development of this brain region.

## ACKNOWLEDGMENTS

We thank Karine Masse for providing us with P2Y1 plasmid. This work was supported by RFBR KOMFI grant 13-04-40194-H (VAL and AGZ). Experiments with P2Y1 were supported by RFBR grant 18-34-00402 (DDK). Experiments on IP and Western blotting were supported by RSCF grant 14-14-00557 (AGZ), confocal imaging was supported by RFBR grant 18-34-00574 (AMN).

## AUTHOR CONTRIBUTIONS

DDK performed the majority of experiments and participated in their planning. VAL conceived and designed the bioinformatic part of the work. VAL and AVS developed the method and algorithm of the lost and gain genes. LIR implemented the programs and carried out the computations. OAZ gathered and analyzed the input data. LIR and AVS analyzed the results. ASI performed experiments on qT-PCR. NYuM performed experiments on IP and Western blotting analysis. AMN conducted experiments with confocal microscopy. MBT performed in situ hybridization and histological sectioning. LP analyzed c-Answer expression at single-cell resolution level. AGZ and VAL generated the general idea of algorithm for search of lost and gain genes. AGZ organized all the work, planned experiments, wrote the text. All authors reviewed and approved the final manuscript.

## MATERIALS AND METHODS

### Bioinformatic analysis, raw score identification, orthologs definition

We define the raw score for any two genes *i* and *j* as the maximum of the BLAST raw scores for all ordered pairs of proteins: one protein for *i* and the other for *j*. The novel score between given genes *m* and *n* is defined by the formula:

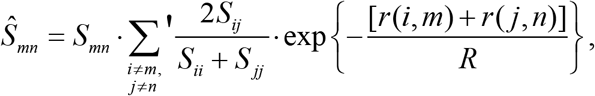

where R is the normalization parameter, and *r* is the distance between the genes *i* and *m, j* and *n*. The summation is carried out over all pairs *i* and *j* of genes in the same contigs (*i* for *m* and *j* for *n*). Moreover, the sum includes not all summands (as indicated by the prime mark).

Sorting all summands in descending order gives us a numeric sequence. The first summand in the equation has the index {*i_0_*, *j_0_*}, now let us remove of members with the indexes *i_0_* or *j_0_* from the sequence; then the second summand is the first member of the resulting sequence, etc. The equation allows that *m=n*.

For the definition (*i*) of homologs in section Results, the BLAST program(Camacho et al., 2009) and the new score are used. Specifically, given *X* gene in the basic species, its homolog *X\** in another species is chosen from genes, for which the E-value corresponding to the BLAST-made alignment of *X* and *X\** is below the specified threshold, so that the raw score is at its maximum (*X\** is the best hit for *X*, BH). In addition, *X* can similarly correspond to *X\** using the same score (*X\** and *X* are bidirectional best hits, BBH). Routinely, BH and BBH are used for upper and lower species, respectively; although other variants are permissible. For the new score the choice of *X\** is postponed until all raw scores have been calculated, which allows calculating the new scores and then acting as above. The definition (*ii*) considers genes as orthologous and paralogous (with no account of synteny) if their proteins belong to the same cluster generated using the new score. Proteins of all considered species are clustered using an original program described in (Zverkov et al., 2012) and extensively approved for the raw score elsewhere (Lyubetsky et al., 2013; Rubanov et al., 2016; Zverkov et al., 2012, 2015). The definition (*iii*) is based on the reconciliation of the gene trees against the species tree, which allows to distinguish duplication and speciation events, thus inferring orthologs and paralogs (also with no account of synteny). This approach was used in the Ensembl Compara database (Vilella et al., 2008) from which we obtain the tables of orthologs and paralogs for the available genes. An original distinction of such genes proposed in (Lyubetsky et al., 2017; Rubanov et al., 2016) can be used as an alternative. Several orthology inference methods including the local synteny consideration were compared on five mammalian genomes in (Jun et al., 2009b).

The new score can take the phylogenetic positions of the considered species into account. Highly conserved elements are important to be considered as witnesses, an algorithm and program for their identification have been proposed in (Rubanov et al., 2016). In a wider context, the gene loss in our approach is considered as a combination of considerable change in nucleotide composition and exon-intron structure as well as significant changes in the related synteny and tissue-specific expression. However, these aspects are ignored in the calculations performed in this study. The implementation of main points of the method can be found at http://lab6.iitp.ru/en/lossgainrsl/. The program is deeply parallelized and can operate on a supercomputer, which is essential if a lot of complete genomes are jointly considered or synteny blocks consist of many neighboring genes. The calculations were carried out on an MVS-10P computer at the Joint Supercomputer Center of the Russian Academy of Sciences.

### Manipulations with tadpoles

Animal experiments were performed in accordance with guidelines approved by the Shemyakin-Ovchinnikov Institute of Bioorganic Chemistry (Moscow, Russia) Animal Committee and handled in accordance with the Animals (Scientific Procedures) Act 1986 and Helsinki Declaration. *X. laevis* tadpoles were obtained, amputated and harvested as we described previously (Ivanova et al., 2013b, 2015). The amputations of *X. laevis* tails were performed using Vannas microscissors after anesthesia with 0,1% tricaine (Sigma-Aldrich).

### DNA constructs, luciferase assay

Cloning strategies are described in Table S1. Luciferase assay was performed as described (Bayramov et al., 2011). Embryos were injected at 2–4 cell stage with synthetic *c-Answer* + *Fgf8* mRNAs or solely *Fgf8* mRNA (3-4 nl of 100ng/μl mRNA water solution per embryo) mixed with one of the luciferase reporter plasmids: AP-1-Luc pGL4.44[luc2P/AP1 RE/Hygro] (Promega) sensitive to the stress-activated MAPK/JNK pathway, or SRE-Luc pGL4.33[luc2P/SRE/Hygro] (Promega) sensitive to MAP/ERK pathway, and the reference pRenilla plasmid (50 pg/embryo of each reporter plasmid). Animal cap explants were excised from the injected embryos at stage 10, cultured until stage 11, selected in three replicate samples by 10 explants in each and processed for luciferase analysis according to Promega protocol.

### Synthetic mRNA and in situ hybridization

Synthetic mRNAs (see Table S1) were prepared with mMessage Machine SP6 Kit (Ambion) after linearization of pCS2-based plasmids with NotI and injected into 2-4 cell stage embryo (3-4 nl of 100ng/μl mRNA water solution per embryo) either into one half of embryo or into the whole embryo or into a particular blastomere in dependence of experiment design. Whole-mount in situ hybridization with antisense probes to *c-Answer, Rx, Pax6, Six3, Engrailed* (Ermakova et al., 2007) was performed as described(Harland, 1991). All mRNAs were mixed with Fluorescein Lysine Dextran (FLD) (Invitrogen, 40 kD, 5 μg/μl) before injections.

### Morpholino oligonucleotides and statistical analysis of malformations in regenerating tadpoles

Morpholino antisense oligonucleotides (GeneTools) to *c-Answer* (*c-Answer* MO) and the control, mismatched, variant of *c-Answer* MO (see Table S1) were injected in water solution with MO final concentration 0.25 mM in volume of 3–4 nl per embryo at 2–4-cell stage. All MOs were mixed with Fluorescein Lysine Dextran (FLD) (Invitrogen, 40 kD, 5 μg/μl) before injections. Embryos were incubated until stage 40-42, at which their tails were inspected using fluorescent stereomicroscope Leica M205 and amputated with micro-scissors (Gills-Vannas scissors). On 7-8 dpa tadpoles with both normally and abnormally regenerated tails were counted. Statistical significance was determined with the paired sample t-test and was set P < 0,01. In sum, 248 tadpoles were analyzed in three independent experiments. Statistical significance was determined with the paired sample t-test, p < 0,001.

### qRT-PCR

Sample preparation and qRT-PCR procedure was performed as described (Ivanova et al., 2013a; Xanthos et al., 2002). Regenerating after amputation tales of tadpoles were harvested on 0,1,2,6 dpa (0 dpa sample was the piece of stump harvested just after amputation considered as the basal control) and were subjected to qRT-PCR with primers to *c-Answer* (see Table S1) and two housekeeping genes, *Ef-1alpha* and *ODC*.

### Immunoprecipitation and antibodies

Lysates of embryos were prepared as described (Bayramov et al., 2011; Martynova et al., 2013). For immunoprecipitation, aliquots of lysates containing standard amount of tagged protein were mixed with EZview Red ANTI-MycAffinity Gel (Sigma E6654) or EZview Red ANTI-Flag Affinity Gel (Sigma F 2426) and incubated with rotation overnight at4C. After washing 5 times with IPB (immunoprecipitation buffer), protein complexes were eluted with 0.15 mg/ml 3xMYC peptide (Sigma) or 3xFLAG peptide (Sigma) accordingly and analyzed by blotting with monoclonal anti-FLAG Alkaline phosphatase conjugated antibodies or anti-MYC Alkaline phosphatase conjugated antibodies accordingly as described previously(Martynova et al., 2008). Myc-/Flag-epitope-containing proteins were produced in *X. laevis* embryos by using constructs in pCS2 plasmids (see Table S1).

### Confocal and fluorescent microscopy, detection of Case12 reporter protein signal

Distribution of EGFP-labeled cells of embryos injected with EGFP-c-Answer mRNA was detected on confocal microscope. All confocal images and FRAP experiments were performed with the confocal microscope “Leica DM IRE 2” using HCX PL APO 63x objective, Ar/Kr laser (488 nm) for excitation of EGFP-tagged proteins. The confocal imaging were obtained at the early-midgastrula stages (stages 10–11) embryos preliminary injected at 4-cell stage with synthetic RNA template (usually 70 pg/blastomere). The *in vivo* fluorescence detection was performed using the fluorescent stereomicroscope Leica M205 and photographed with Leica camera (DC 400F). To test effects of c-Answer on this signaling through P2Y1 receptor, embryos were injected with mRNA of Ca^2+^-sensitive variant of green fluorescent protein (Case12 reporter protein) mixed with *P2Y1* and *c-Answer* mRNAs or solely *P2Y1* mRNA and incubated until stage 11. Animal cap’s explants were excised from the injected embryos and dissociated on single cells in Calcium Magnesium Free Medium (CMF) (116 mM NaCl, 0.67 mM KCl, 4.6 mM Tris, 0.4 mM EDTA). Ca^2+^ influx upon the addition of P2Y1 agonist ADP (Sigma) to 300 mkM final concentration was recorded via measuring the fluorescence of Case12 reporter on the fluorescent stereomicroscope Leica M205 with Leica camera (DC 400F). The maximal value of signal produced after adding of ADP in each round of measurement was used for further statistical analysis. 40 embryos were analyzed in three independent experiments for each variant of injected mRNA. Statistical significance was determined with the paired sample t-test, p ≤ 0,01.

### Cryosectioning

After fixation in 4% PFA the *X. laevis* tail and hind limb bud samples were transferred to the melted warm (+47°C) 1,5% bacto-agar on 5% sucrose solution and were oriented until the sample curdled. The cube with the sample was left in 30% sucrose solution for 12-15 hours and then was bound by the Neg-50 (Richard-Allan Scientific) to the specimen holder and covered by Neg-50. Further, the holder with the sample was carefully inserted into a liquid nitrogen and then cryosectioned (20μ thick) on the Microm HM 525 (Thermo Scientific) and placed on superfrost plus microscope slides (Fisher Scientific, cat.# 12-550-15). Cell nuclei were then stained on these sections with DAPI solution.

### CRISPR/Cas9 knockout and genotyping

Target site for the 2^nd^ exon of *c-Answer* was designed in http://chopchop.cbu.uib.no/software (see Table S1). sgRNA template construction and template assembly by PCR was performed as described (Nakayama et al., 2014). In vitro transcription of sgRNA was carried out with kit GeneArtTM PlatinumTM Cas9 Nuclease Ready-to-transfect wild-type Cas9 protein for performing CRISPR/Cas9-mediated genome editing Catalog Numbers B25640, B25641 (Invitrogen) according to the guidelines enclosed. Cas9 protein from the kit was mixed with *c-Answer* sgRNA with final concentration 0.8ng and 400pg accordingly in water solution with Fluorescein Lysine Dextran (FLD) (Invitrogen, 40 kD, 5 μg/μl), incubated for 5 min at RT and injected into embryos 20min after fertilization. Embryos were incubated until stage 12-14 and then total DNA was extracted from 10 randomly selected embryos as described(Sive et al., 2010) for method validation. The cDNA of the region of putative mutation was obtained by PCR with direct and reverse specific exterior primers flanking the mutated region of *c-Answer*. 50ng of the obtained PCR product was taken for another PCR (total volume 25 μl, 40 cycles) with the interior direct primer (1 μl of 100 pmol/μl), reverse adapter primer (reverse interior primer with the adapter part common for all barcode primers) (1μl of 10 pmol/μl) and bar-code primer unique for each embryo (1μl of 100 pmol/μl. For all primer sequences see Table S1. The obtained PCR products with unique bar-codes for each embryo were mixed together and genotyped by NGS (Illumina MiSeq).

### Single-cell data description

At stage 16 *c-Answer* is expressed by 7% (1004 out of 13478) cells in range of 1-25 mRNA molecules per cell. At stage 18 *c-Answer* is expressed by 11% (1442 out of 12432) cells in range of 1-17 mRNA molecules per cell. At stage 22 *c-Answer* is expressed by 8% (3257 out of 37749) cells in range of 1-20 mRNA molecules per cell. We considered tissues to be enriched *c-Answer* if their cells expressing *c-Answer* give more than 15% of all the cells belonging to the given tissue sub-type. All the tissue sub-types for these stages may be found on: https://kleintools.hms.harvard.edu/tools/springViewer_1_6_dev.html?cgibin/client_datasets/xenopus_embryo_timecourse_v2/cAnswerSt16 for stage 16; https://kleintools.hms.harvard.edu/tools/springViewer_1_6_dev.html?cgibin/client_datasets/xenopus_embryo_timecourse_v2/cAnswerSt18 for stage 18; https://kleintools.hms.harvard.edu/tools/springViewer_1005F;6005F;dev.html?cgibin/client005F;datasets/xenopusembryo005F;timecourse005F;v2/cAnswerSt22 for stage 22; in order to obtain the percentage of cells per tissue sub-type expressing *c-Answer*, gene name should be chosen as LOC100135223 and all the cells expressing it in the range of 1-max UMI per cell (Slider select 1-max) should be laid out (Layout, Move left). Then the cells expressing *c-Answer* should be positive selected and option Celltype should be turned on.

## SUPPLEMENTARY MATERIALS

### Species used to identify lost genes

The used species are well represented in the Ensembl database (Zerbino et al., 2018). The assembly name is specified for each genome below (with the GenBank accession number in parentheses if available). <u>Fish</u>: agnathous fish – *Petromyzon marinus* (lamprey) Pmarinus_7.0 (GCA_000148955.1); gnathostomatous fish – *Astyanax mexicanus* (cave fish) AstMex102 (GCA_000372685.1), *Danio rerio* (zebrafish) GRCz10 (GCA_000002035.3), *Gasterosteus aculeatus* (stickleback) BROAD S1 (GCA_000180675.1), *Latimeria chalumnae* (coelacanth) LatCha1 (GCA_000225785.1), *Lepisosteus oculatus* (spotted gar) LepOcu1 (GCA_000242695.1), *Oreochromis niloticus* (tilapia) Orenil1.0 (GCA_000188235.1), *Oryzias latipes* (medaka) HdrR, *Poecilia formosa* (amazon molly) Poecilia_formosa-5.1.2 (GCA_000485575.1), *Takifugu rubrípes* (fugu) FUGU 4.0 (GCF_000180615.1, FUGU5), *Tetraodon nigroviridis* (tetraodon) TETRAODON 8.0 (GCA_000180735.1), *Xiphophorus maculatus* (platyfish) Xipmac4.4.2 (GCA_000241075.1); <u>amphibians</u>: *Xenopus tropicalis* (clawed frog) JGI_4.2 (GCA_000004195.1); <u>reptiles</u>: *Anolis carolinensis* (anole lizard) AnoCar2.0 (GCA_000090745.1), *Pelodiscus sinensis* (chinese softshell turtle) PelSin_1.0 (GCA_000230535.1); <u>birds</u>: *Gallus gallus* (chicken) Gallus_gallus-5.0 (GCA_000002315.3), *Meleagris gallopavo* (turkey) Turkey_2.01 (GCA_000146605.1), *Taeniopygia guttata* (zebra finch) taeGut3.2.4 (GCA_000151805.1), *Anas platyrhynchos* (duck) BGI_duck_1.0 (GCA_000355885.1), *Ficedula albicollis* (flycatcher) FicAlb_1.4 (GCA_000247815.1); <u>primitive mammals</u> – *Ornithorhynchus anatinus* (platypus) OANA5 (GCF_000002275.2), *Sarcophilus harrisii* (Tasmanian devil) DEVIL7.0 (GCA_000189315.1), *Monodelphis domestica* (opossum) monDom5 (GCF_000002295.2), and <u>placental mammals</u> – *Homo sapiens* (human) GRCh38.p10 (GCA_000001405.25), *Mus musculus* (mouse) GRCm38.p5 (GCA_000001635.7), and *Cavia porcellus* (Guinea pig) cavPor3 (GCF_000151735.1). Our algorithm predicts the same set of lost genes for more recent GenBank sequencing data.

**S1 Table Supplementary Table S1.**
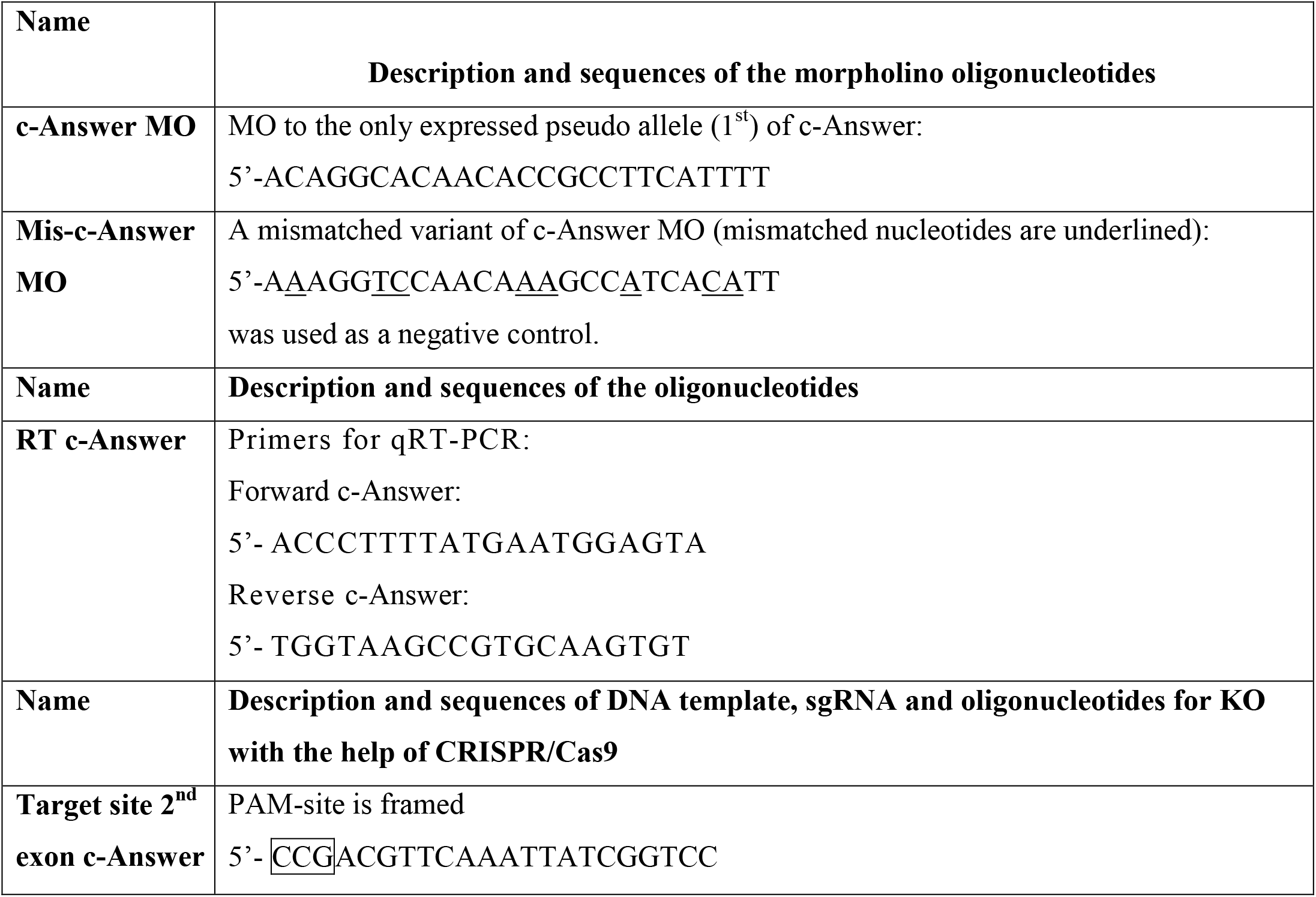
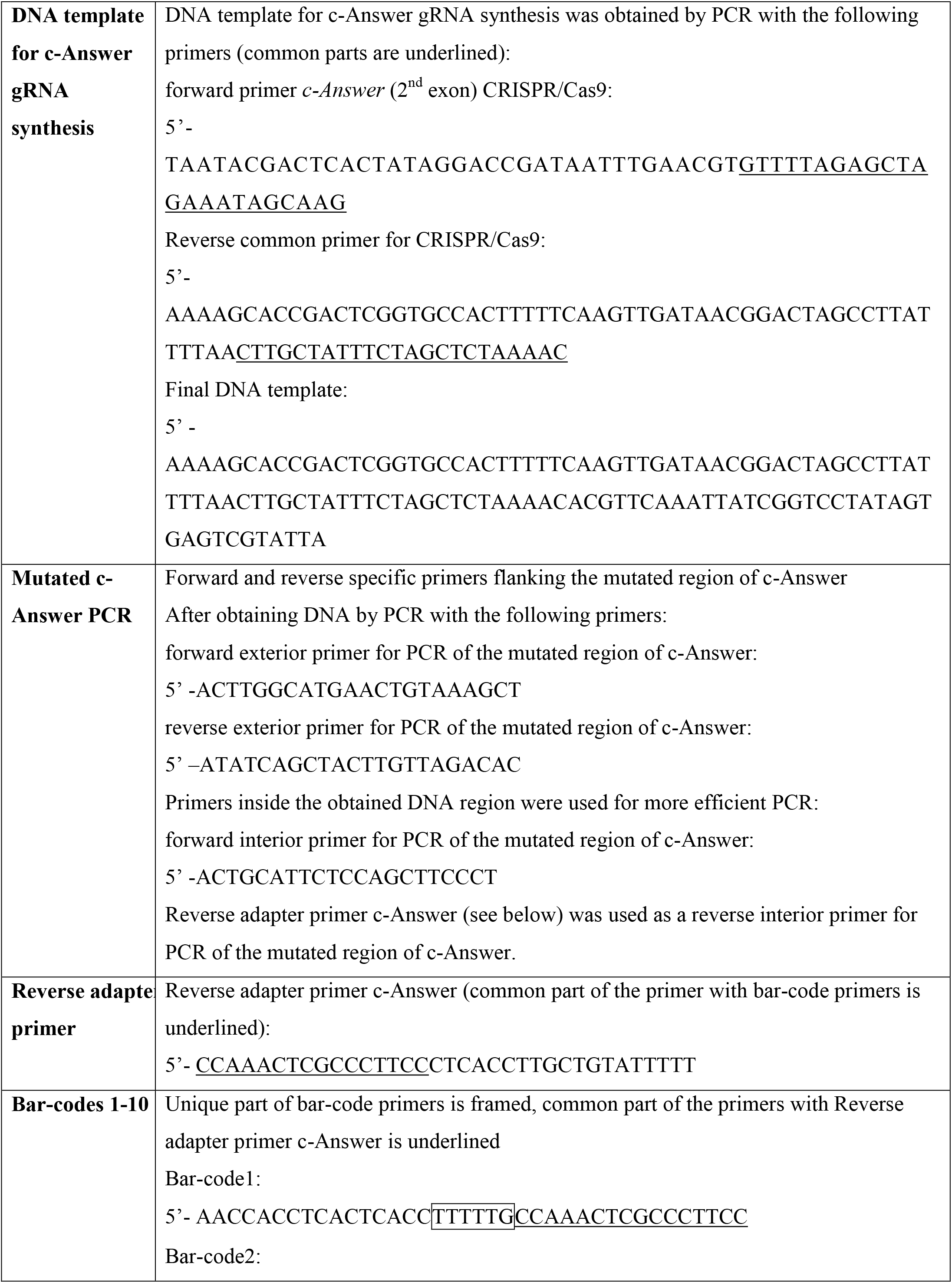
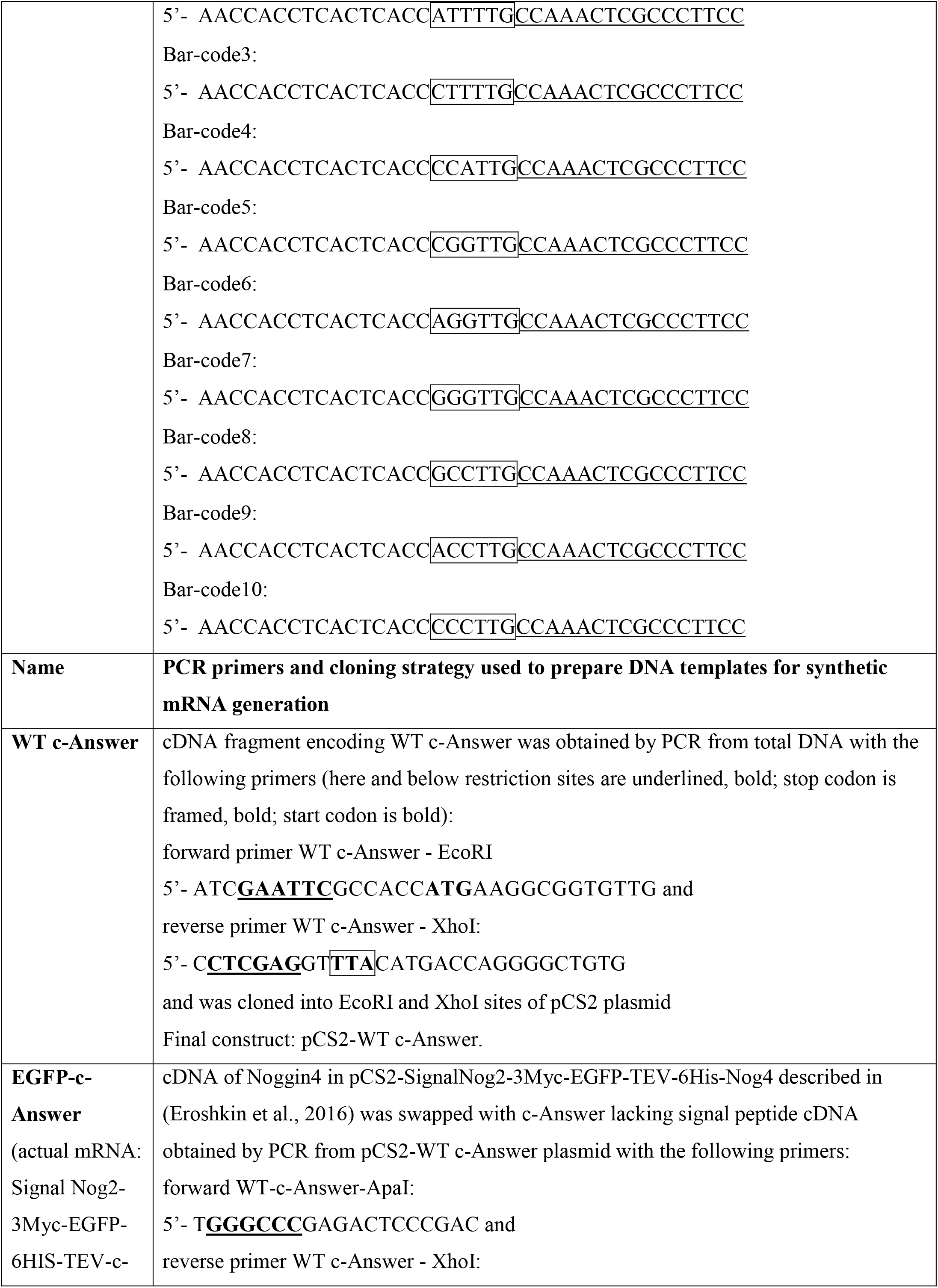
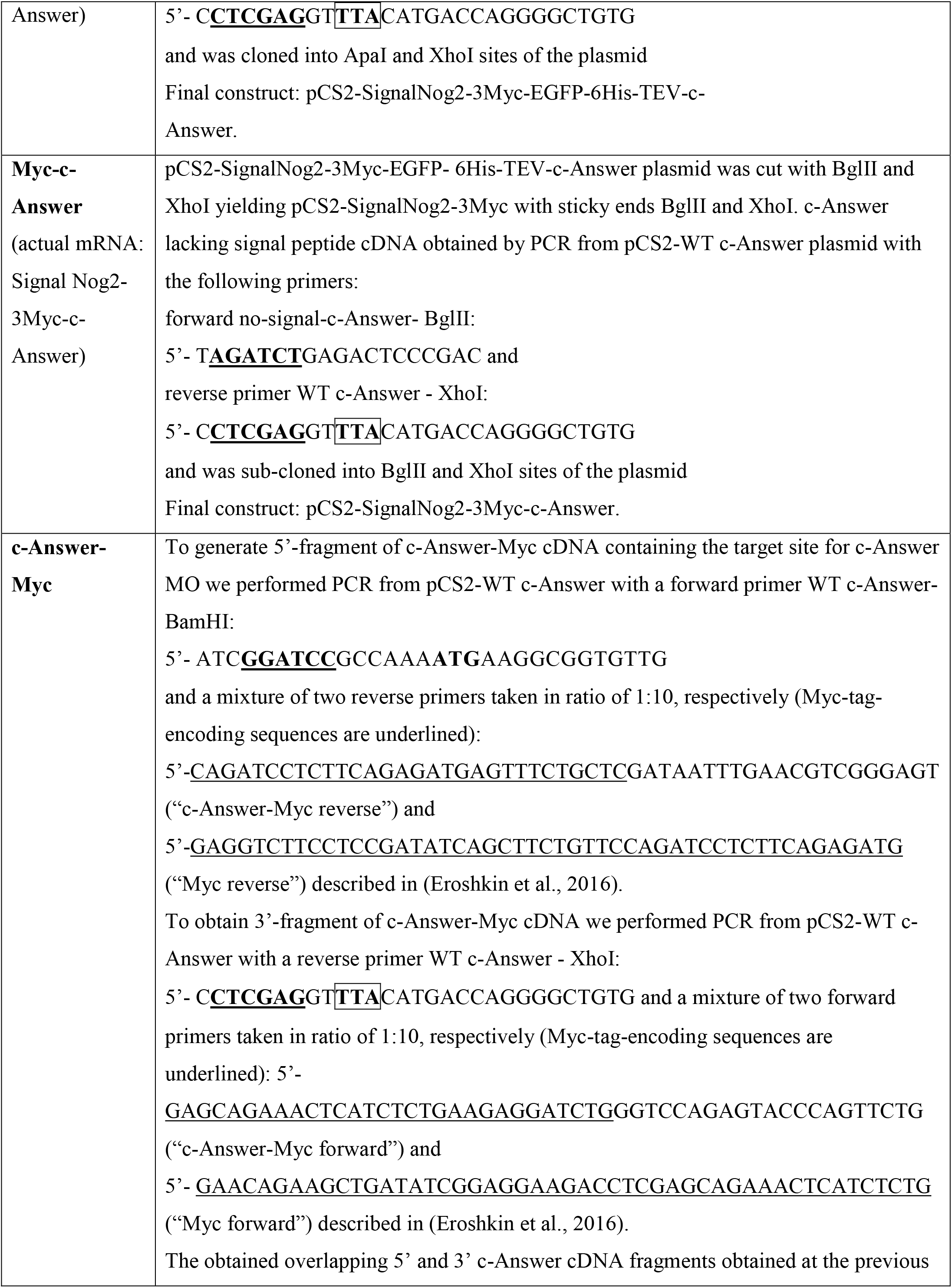
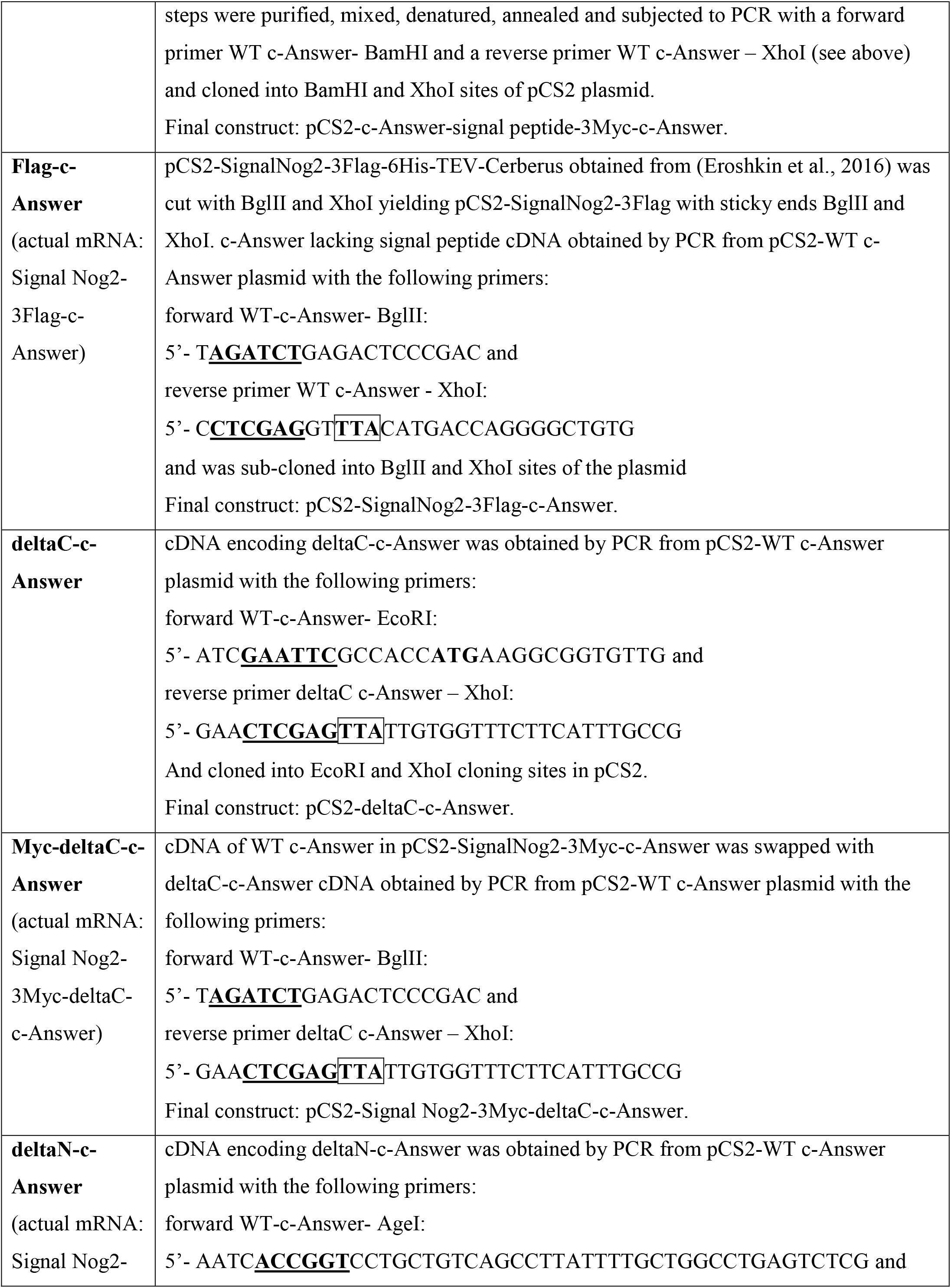
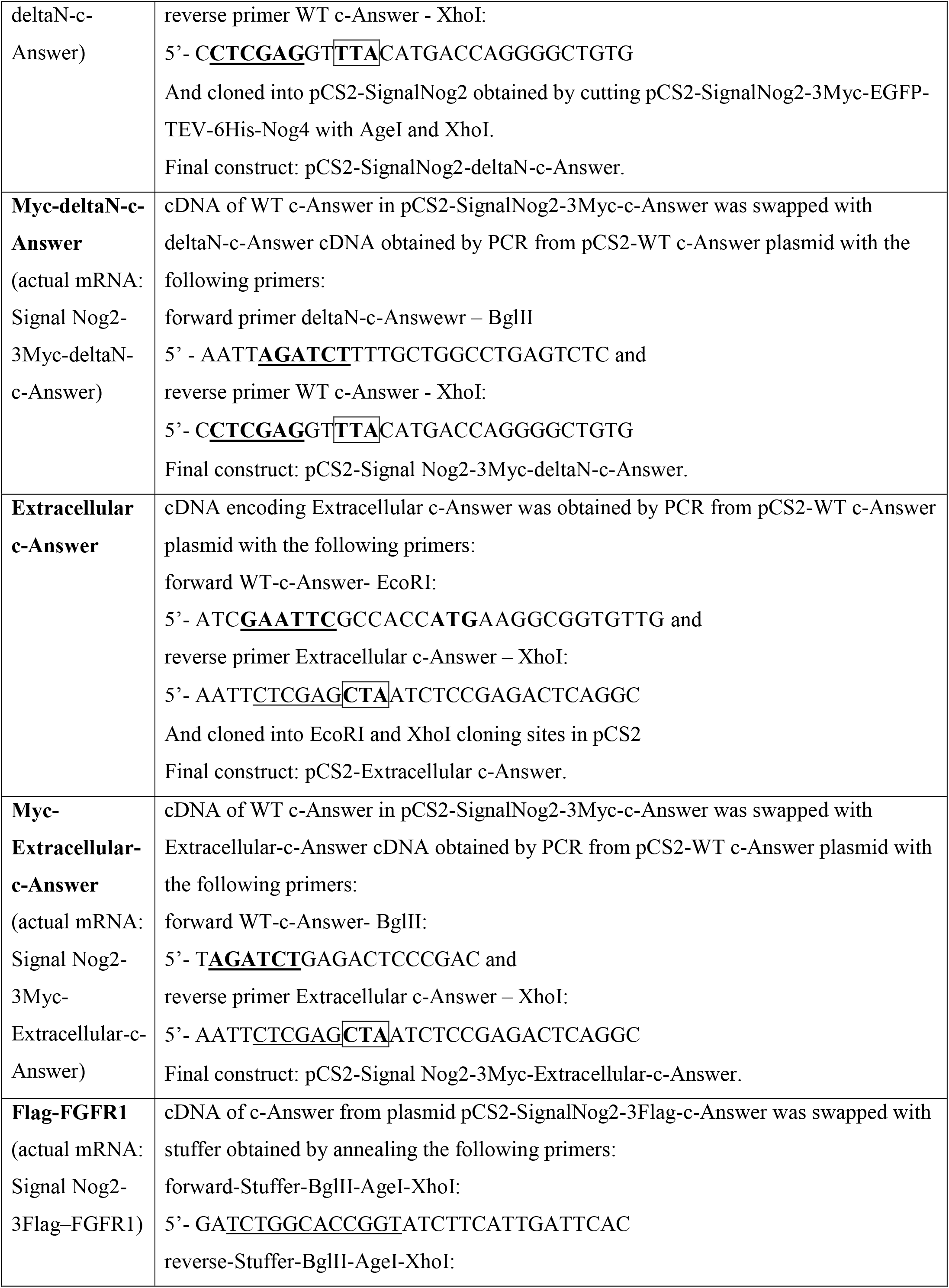
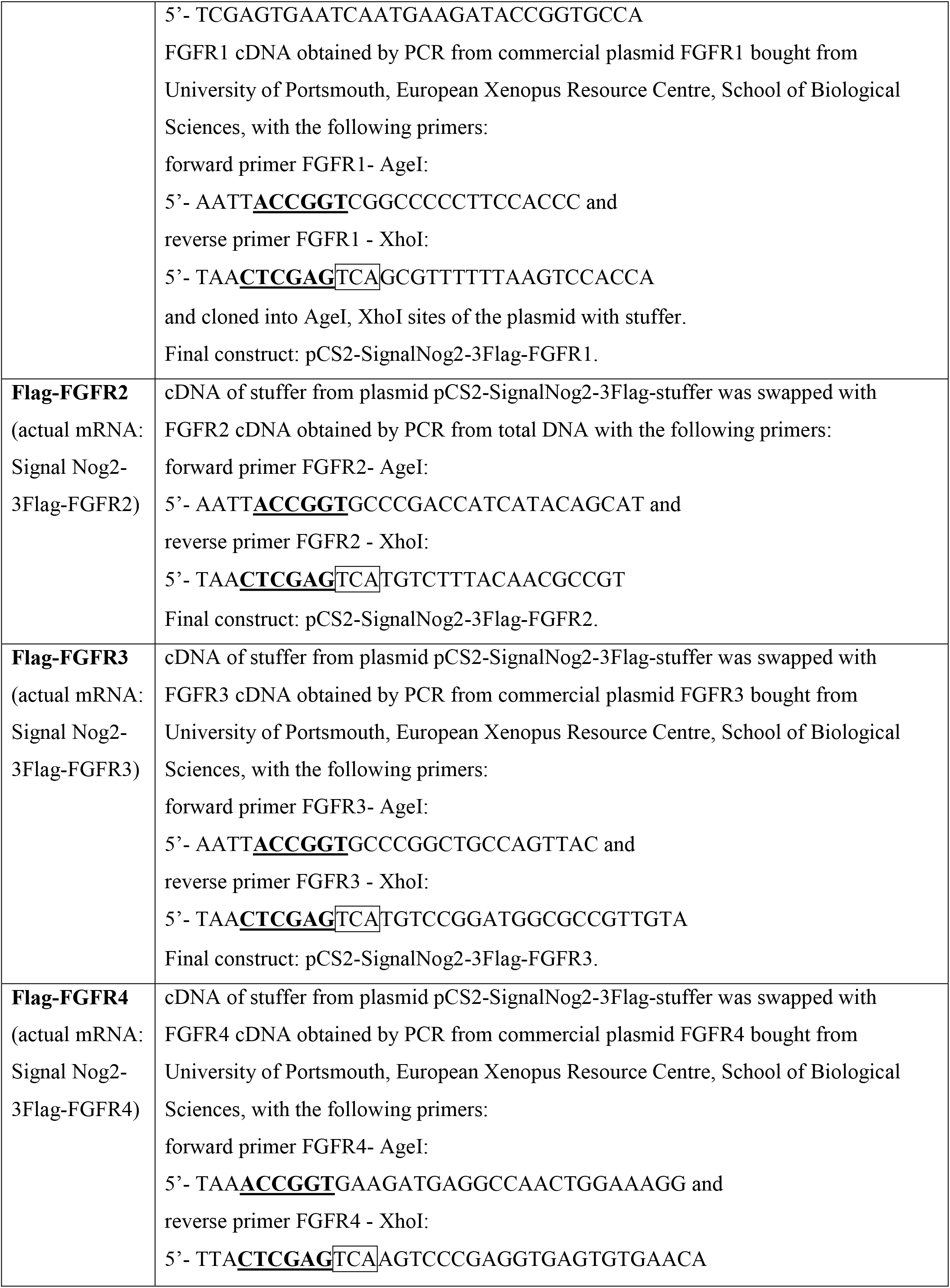
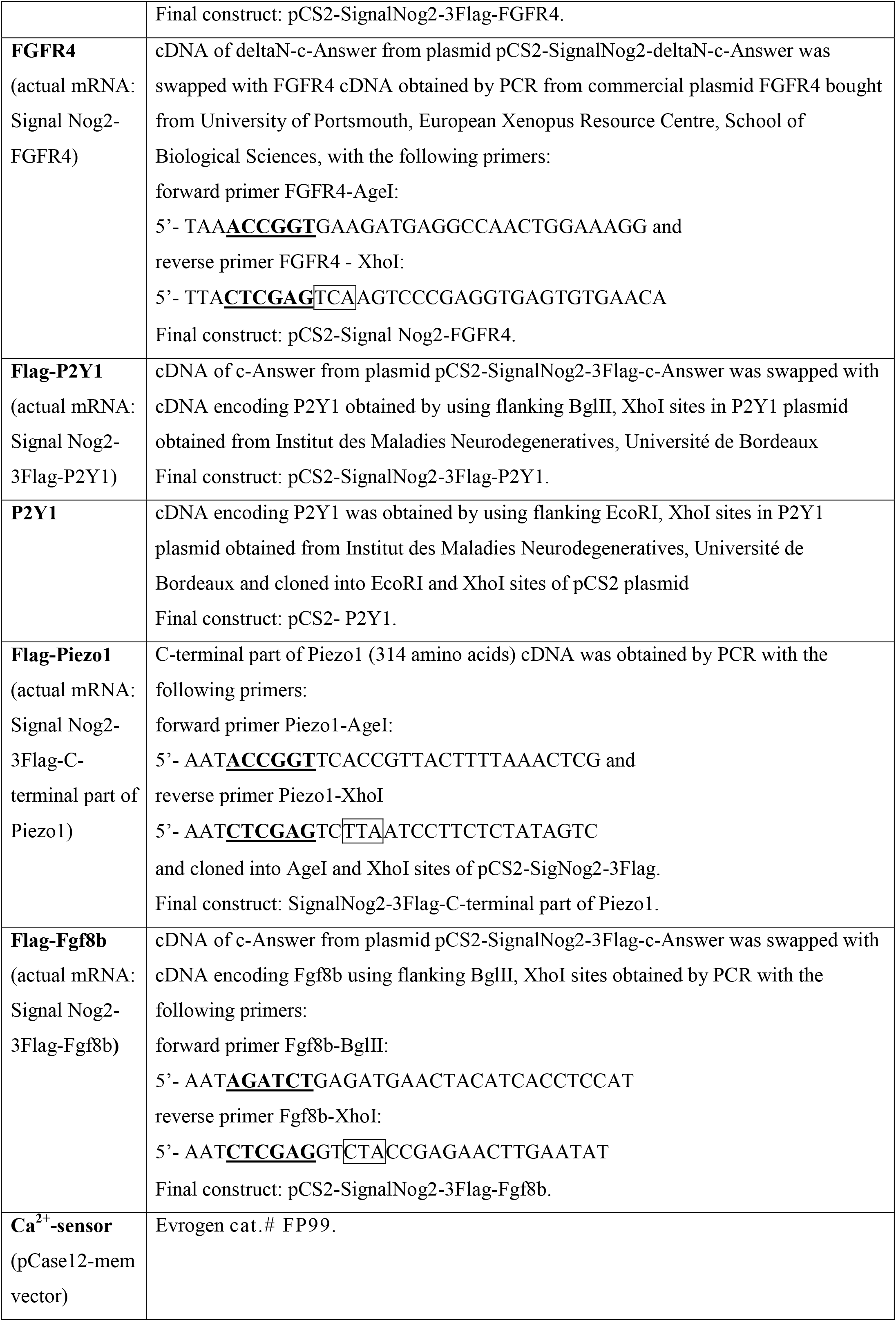
All DNA constructs were checked by sequencing.

### Supplementary Figures

**Figure S1.**
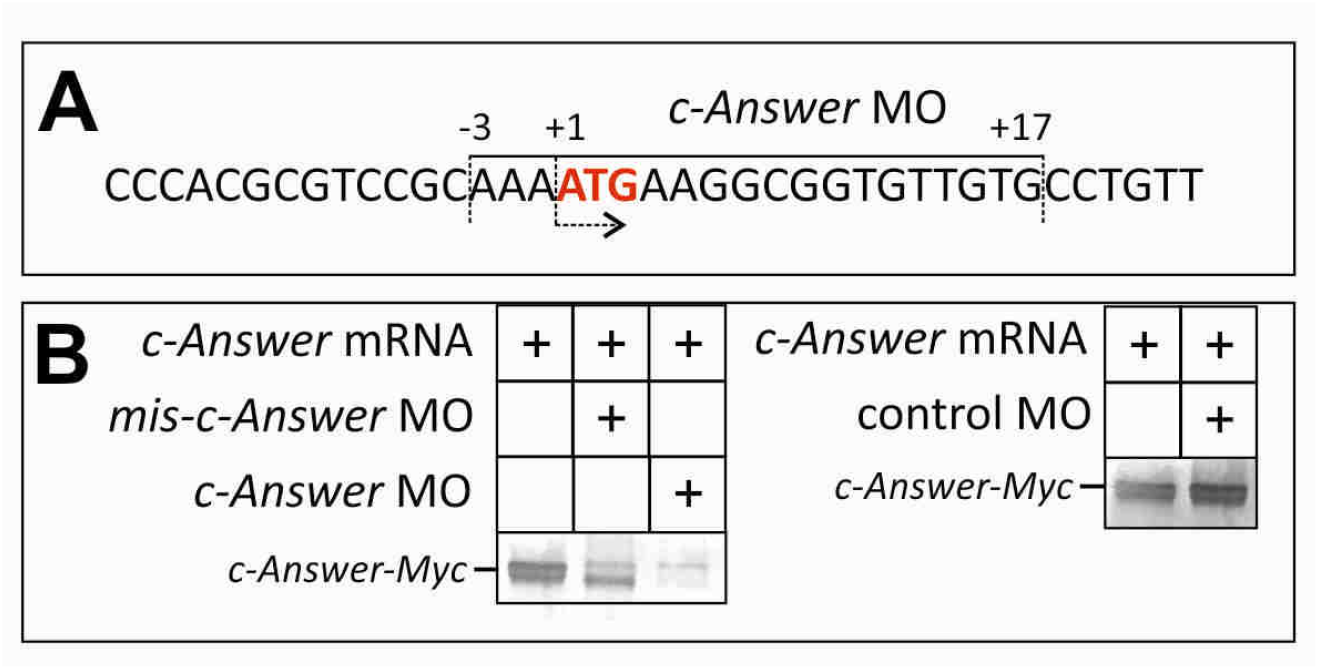
Testing of MO specificity and efficiency. **A**. Scheme of the morpholino target site location on pseudoallele A of the *X. laevis c-Answer* mRNA. **B**. *c-Answer-Myc* mRNA was injected into each blastomere of 2-cell *X. laevis* embryos (100pg/blastomere) either alone or in mixture with control *mis-c-Answer* MO (8nl of 0.2 mM water solution) or the standard control MO provided by GeneTools (8 nl of 0.5 mM water solution). The injected embryos were collected at the middle gastrula stage and analyzed for presence of c-Answer-Myc by Western blotting with anti-Myc antibody (see Materials and Methods for details).

**Figure S2.**
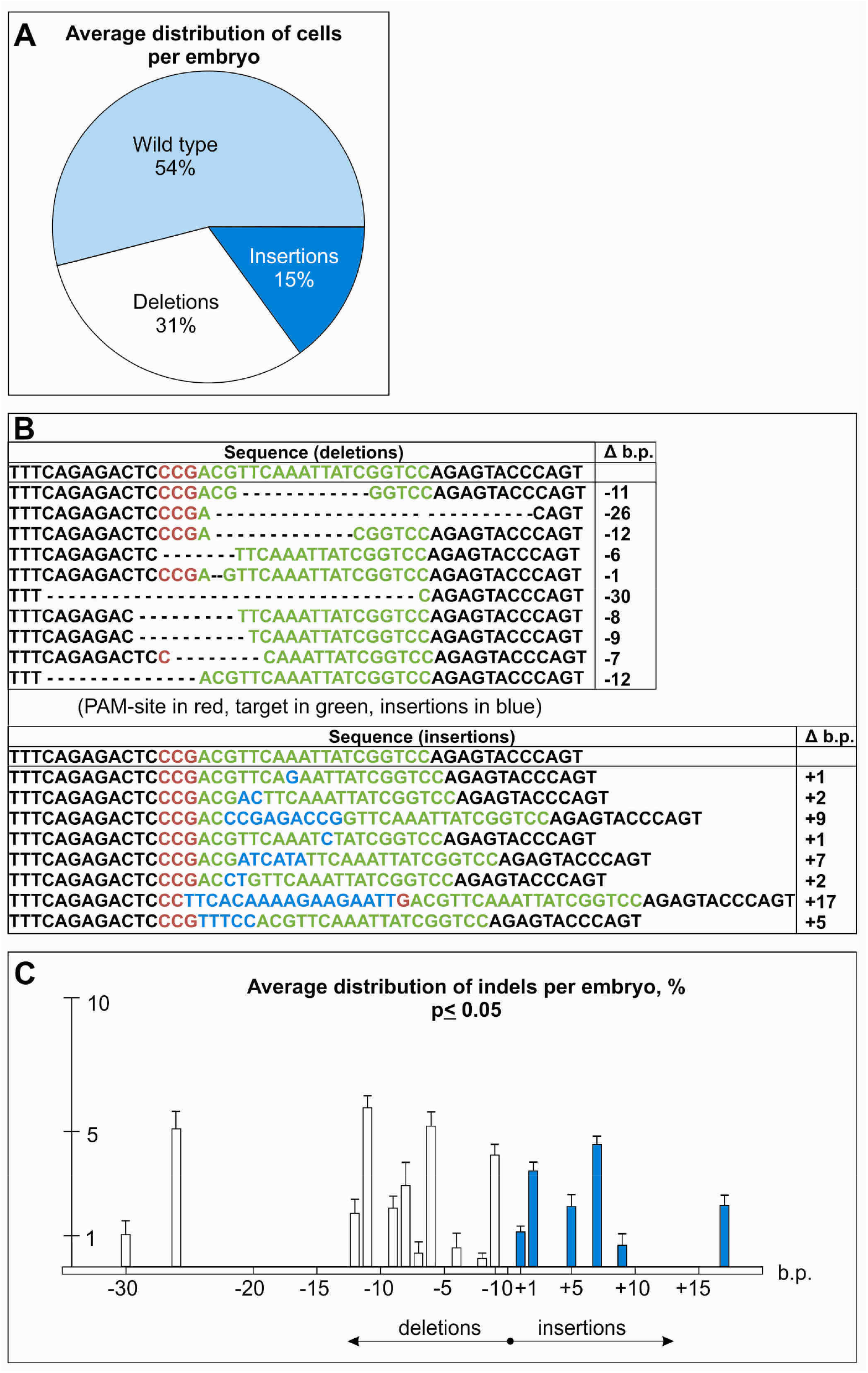
The results of embryo genotyping after c-Answer CRISPR/Cas9 knockout. **A**. Diagram showing the average distribution of cells with insertions/deletions/no mutations per embryo with *c-Answer* CRISPR/Cas9 knockout (data obtained from 10 embryos). **B**. The most frequent variants of insertions and deletions (data summarized from 10 embryos). **C**. Diagram showing the average frequency of insertions and deletions of the indicated length (b.p.) per embryo, data is averaged from 10 embryos.

**Figure S3.**
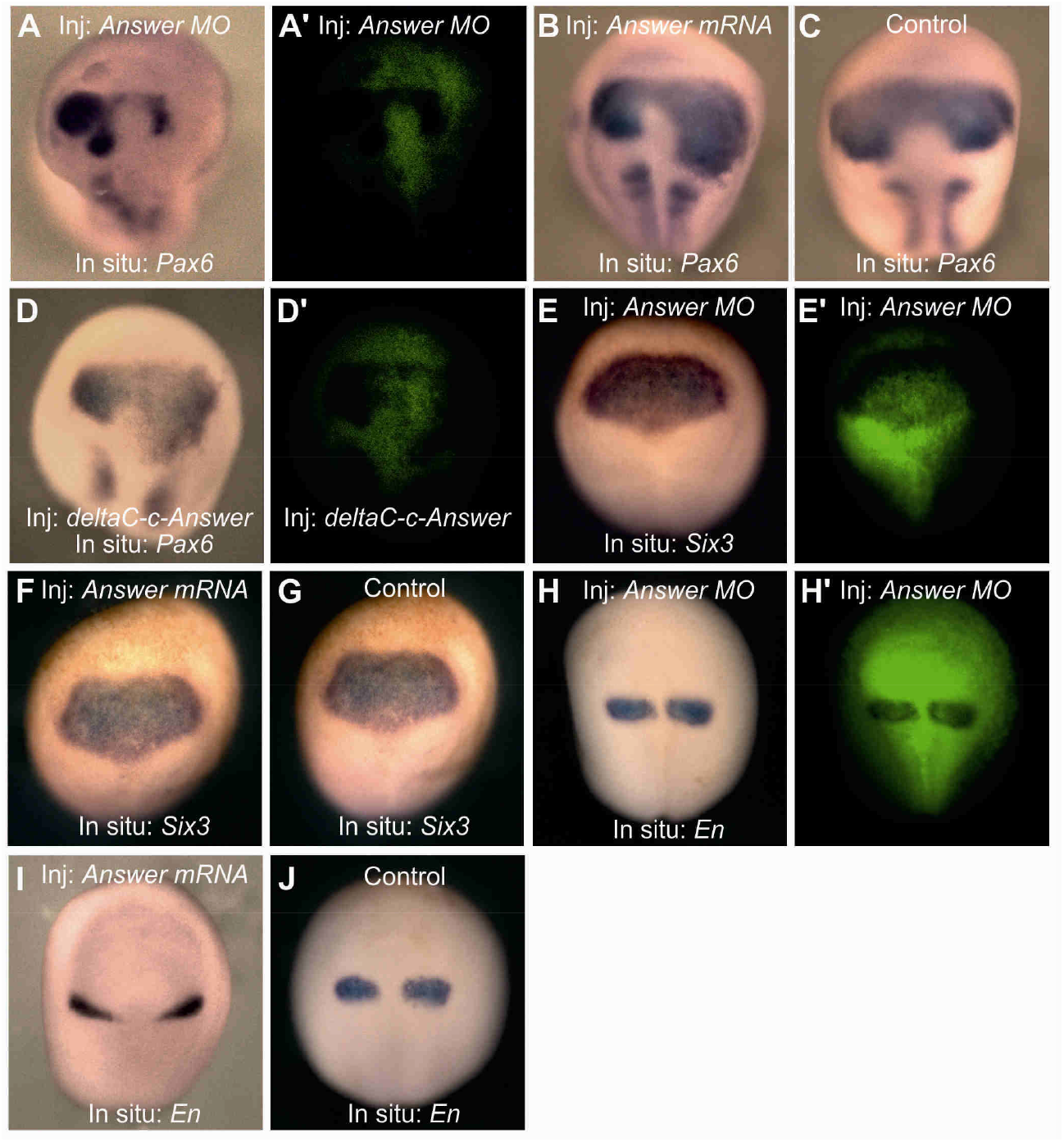
Whole-mount in situ hybridization with probes to the transcripts of the indicated genes of embryos injected with *c-Answer* MO or *c-Answer* mRNA variants. **A**. Inhibition of the *Pax6* expression on the right side of the middle neurula embryos injected with *c-Answer* MO into the right dorsal blastomere at 4-cell stage. Left side is not injected control. **A’**. The fluorescent image for **A** demonstrating the distribution of green fluorescent tracer, FLD, coinjected with *c-Answer* MO. **B**. Ectopic *Pax6* expression in the middle neurula embryos injected with *c-Answer* MO into 2 blastomeres at 2-cell stage. **C**. *Pax6* expression in the control (not injected) middle neurula embryos. **D**. Ectopic *Pax6* expression on the right side of the middle neurula embryos injected with deltaC-*c-Answer* mRNA into the right dorsal blastomere at 4-cell stage. Left side is not injected control. **D’**. The fluorescent image for **D** demonstrating the distribution of green fluorescent tracer, FLD, coinjected with *deltaC-c-Answer* mRNA. **E**. No effect on *Six3* expression in the middle neurula embryos injected with *c-Answer* MO into the right dorsal blastomere at 4-cell stage. **E’**. The fluorescent image for **E** demonstrating the distribution of green fluorescent tracer, FLD, coinjected with *c-Answer* MO. **F**. No effect on *Six3* expression in the middle neurula embryos injected with *c-Answer* mRNA into 2 blastomeres at 2-cell stage. **G**. *Six3* expression in the control (not injected) middle neurula embryo. **H**. No effect on *En* expression in the middle neurula embryos injected with *c-Answer* MO into 2 blastomeres at 2-cell stage. **J** can be referred to as not injected control. **H’**. The fluorescent image for **H** demonstrating the distribution of green fluorescent tracer, FLD, coinjected with *c-Answer* MO. **I**. Inhibition of the *En* expression in the middle neurula embryos injected with *c-Answer* mRNA into 2 blastomeres at 2-cell stage. **J**. *En* expression in the control (not injected) middle neurula embryo.

**Figure S4.**
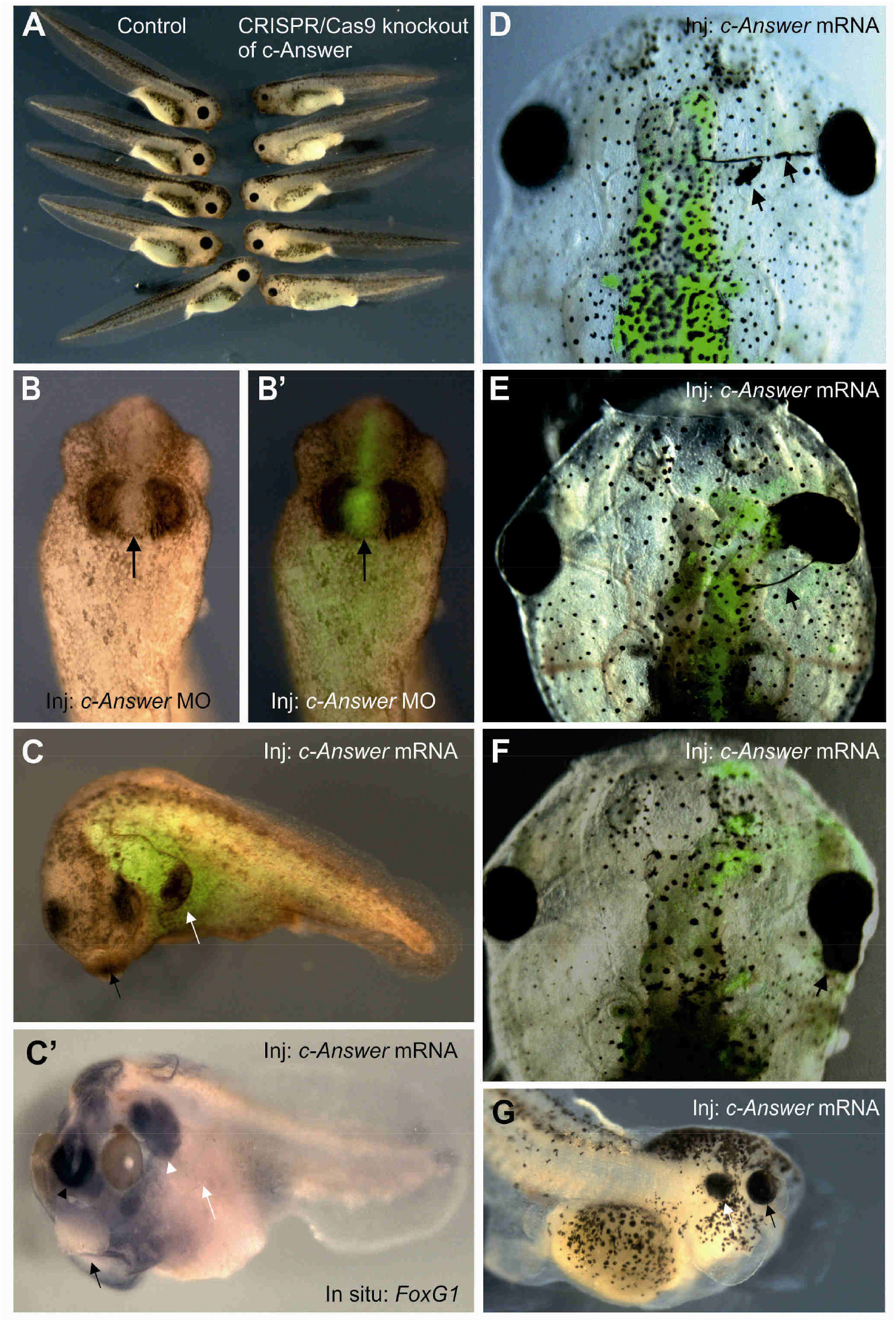
Phenotypes of tadpoles after c-Answer knockout and overexpression of c-Answer variants. **A**. Tadpoles with *c-Answer* knockout (right) have smaller eyes than the control ones (left). **B and B’**. Local injection of c-Answer MO mixed with FLD tracer in one blastomere, which give rise to the middle part of the cement gland, at 16-cell stage resulted in the inhibition of the cement gland differentiation in this part. The embryo at stage 26 is shown from the ventral side, anterior to the top. **C**. Tadpole overexpressing wild-type c-Answer has ectopic cement gland on the head process (arrow). **C’**. Whole-mount in situ hybridization with the probe to *FoxG1* of the same tadpole as on B reveals additional telencephalon within the head process (arrow head). **G**. Ectopic eye in tadpole overexpressing wild-type c-Answer. **D-F**. Ectopic RPE differentiation in tadpoles overexpressing wild-type c-Answer (arrows).

**Figure S5.**
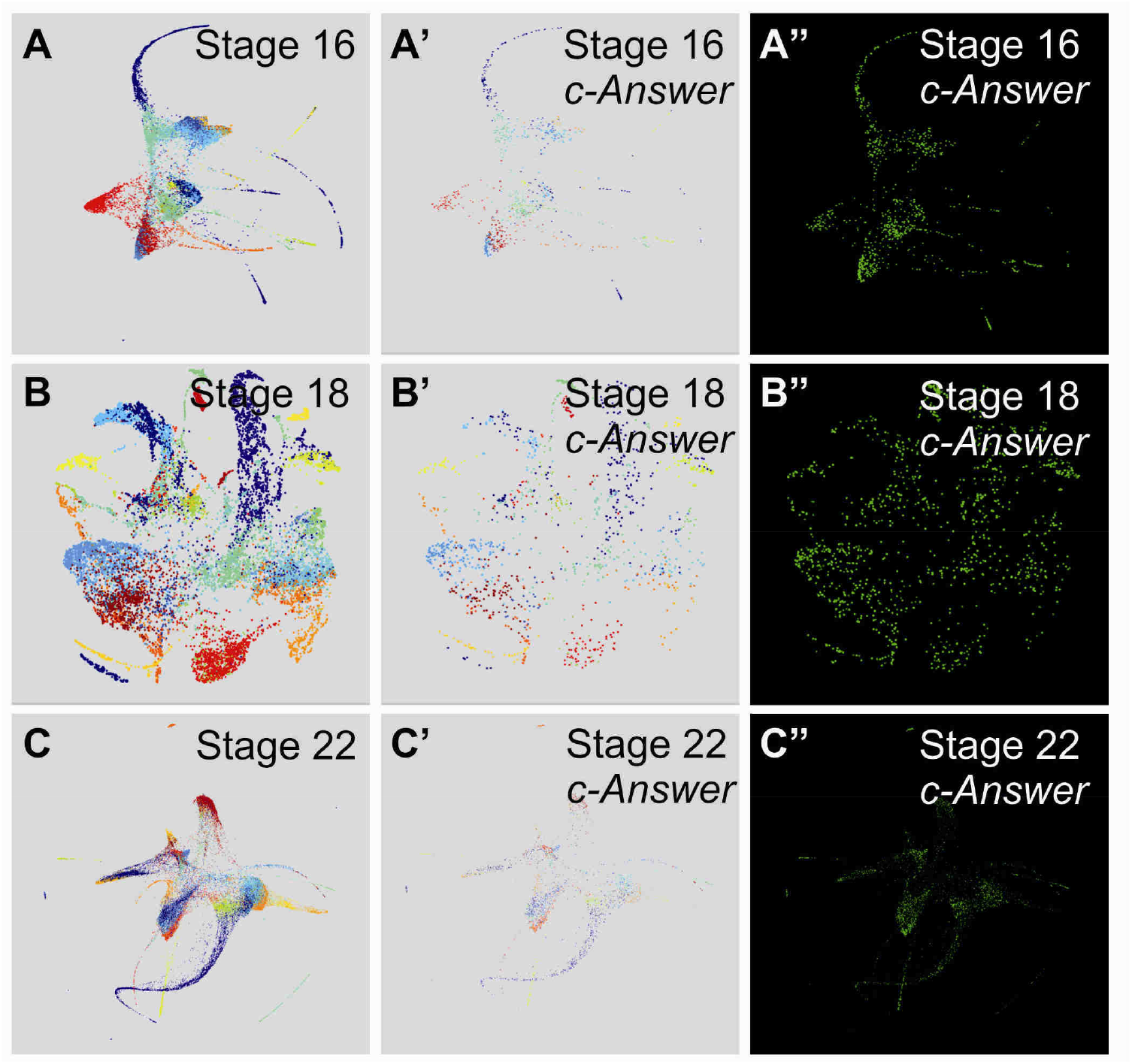
Distribution of c-Answer expression in embryos at stages 16, 18, and 22. **A**. Stage 16, all 13478 embryonic cells arranged by their expression profile similarity (Briggs et al., 2018), clustered and colored according to the tissue subtype, where they belong. For legend see https://kleintools.hms.harvard.edu/tools/springViewer_1_6_dev.html?cgibin/client_datasets/xenopus_embryo_timecourse_v2/cAnswerSt16 **A’**. Stage 16, 1004 cells expressing *c-Answer* are clustered and colored according to the tissue subtype, where they belong. **A”**. Stage 16, 1004 cells expressing *c-Answer* are colored green. The intensity of green color depends on the number of *c-Answer* mRNA present in cell, more intense color relates to greater amount of *c-Answer* mRNA molecules per cell. **B**. Stage 18, all 12432 embryonic cells are clustered and colored according to the tissue subtype, where they belong. For legend see https://kleintools.hms.harvard.edu/tools/springViewer_1_6_dev.html?cgi-bin/clientdatasets/xenopus_embryo_timecourse_v2/cAnswerSt18 **B’**. Stage 18, 1442 cells expressing *c-Answer* are clustered and colored according to the tissue subtype, where they belong. **B”**. Stage 18, 1442 cells expressing *c-Answer* are colored green. The intensity of green color depends on the number of *c-Answer* mRNA present in cell, more intense color relates to greater amount of *c-Answer* mRNA molecules per cell. **C**. Stage 22, all 37749 embryonic cells are clustered and colored according to the tissue subtype, where they belong. For legend see https://kleintools.hms.harvard.edu/tools/springViewer_1_6_dev.html?cgi-bin/client_datasets/xenopus_embryo_timecourse_v2/cAnswerSt22 **C’**. Stage 22, 3257 cells expressing *c-Answer* are colored according to the tissue subtype, where they belong. **C”**. Stage 22, 3257 cells expressing *c-Answer* are colored green. The intensity of green color depends on the number of *c-Answer* mRNA present in cell, more intense color relates to greater amount of *c-Answer* mRNA molecules per cell.

**Video S1. Activation of Ca^2+^ sensor upon ADP addition in dissociated cells of embryos injected with *P2Y1* or *P2Y1+c-Answer* mRNAs (see attached file Video S1)**

Videos in green channel show the fluorescent signal of Case9 reporter in dissociated animal cap cells upon Ca^2+^ flux in the cytoplasm of these cells after ADP application. Red channel is a control showing the distribution of the injected material.

Left-side videos demonstrate cells from embryos injected with *P2Y1* mRNA.

Right-side videos demonstrate cells from embryos injected with *P2Y1+c-Answer* mRNA. The activation of Case9 reporter is higher in cells from embryos injected with *P2Y1+c-Answer* mRNA.

**Video S2. Activation of Ca^2+^ sensor upon ADP addition in whole embryos injected with *P2Y1* or *P2Y1+c-Answer* mRNAs (see attached file Video S2)**

Videos in green channel show the fluorescent signal of Case9 reporter in whole embryos upon Ca^2+^ flux in the cytoplasm of embryo’s cells after ADP application. Red channel is a control showing the distribution of the injected material.

Left-side videos demonstrate embryos injected with *P2Y1* mRNA.

Right-side videos demonstrate embryos injected with *P2Y1+c-Answer* mRNA. The activation of Case9 reporter is higher in embryos injected with *P2Y1+c-Answer* mRNA.

